# A novel framework for expanding RNNs with biophysical detail to solve cognitive tasks

**DOI:** 10.64898/2026.03.13.711746

**Authors:** Nicholas Tolley, Stephanie Jones

## Abstract

Recurrent neural networks (RNNs) have proven to be highly successful in emulating human-like cognitive functions such as working memory. In recent years, RNNs are evolving to incorporate more biophysical realism to produce more plausible predictions on how cognitive tasks are solved in real neural circuits. However, there are major challenges in constructing and training networks with the complex and nonlinear properties of real neurons. A major component of the success of RNNs is that they share the same mathematical base as deep neural networks, permitting highly efficient optimization of model parameters using standard deep learning techniques. To do so, they use abstract representations of neurons which fail to capture the impact of cell-level biophysical and morphologic properties that may benefit network-level function. Expanding task-trained RNNs with biophysical properties such as dendrites and active ionic currents poses substantial challenges, as it moves these models away from the validated training regimes known to be highly effective for RNNs.

To address this gap, we developed a biophysically detailed reservoir computing (BRC) framework with the goal of extracting mechanistic insights from biophysical neural models, and propose that these insights can be used to guide model choices that will work for specific categories of cognitive tasks. The BRC network was constructed with synaptically coupled excitatory and inhibitory cells, in which the excitatory cells include multicompartment biophysically active dendrites; motivated by empirical studies suggesting dendrites have desirable computational benefits (e.g. pattern classification and coincidence detection). We trained the BRC network to do a simplified working memory task where it had to maintain the representation of an extrinsic “cue” input. We studied the impact of extrinsic input time constants (fast AMPA vs slow NMDA) and location (dendrite vs soma) on the ability of a network to solve the task. Our results revealed that cue inputs through NMDA receptors are particularly efficient for solving the working memory task. Further, the properties of NMDA receptors are uniquely suited for cue inputs delivered at the dendrite, as networks trained with dendritic AMPA cue inputs failed to solve the task. Detailed examination of the cell and network dynamics that solve the task reveals distinct local network configurations and computing principles for the different types of extrinsic input. Overall, much like the body of mechanistic insights that have underpinned the success of training RNNs, this study lays the groundwork for applying the BRC framework to train biophysically detailed neural models to solve complex human-like cognitive tasks.

## Introduction

Recurrent neural networks (RNNs) are a highly popular deep learning architecture that have most prominently been used for forecasting and classification tasks involving time series [1]. In recent years, they have been adapted to solve more neuroscience-relevant cognitive tasks that emulate human behaviors, such as working memory [2–5]. Task-trained neural models such as RNNs are useful tools for emulating neural dynamics that can solve cognitive tasks, but they lack biological realism that may convey useful computational properties [6]. While adding biology-inspired properties is a valuable goal for advancing the field, it is important to recognize that the historical success of task-trained neural models is largely due to their shared building blocks with deep neural networks, and the decades of research allowing scientists to know “what works” for training these simplified non-biophysical networks. Deep neural networks don’t work out of the box, their success builds upon the mechanistic insights developed over vast prior research which inform choices such as network architecture [7], data augmentation [8, 9], weight initialization and parameterization [10–12], loss function construction [13, 14], and many other features. When adding biological realism, we are moving entirely away from the validated regimes where we confidently know which choices work, as evidenced by the highly distinct set of methods that have been developed for the training of spiking neural networks (SNN, [15–20]). SNNs already represent a dramatic departure from the basic foundations of RNNs (communication by spikes), however they represent a relatively minor modification with respect to the complexity of biophysically-detailed neural models. These issues highlight a need to develop mechanistic insights that help anticipate which choices work for a given task, and for the level of biological detail added to the network. Establishing mechanistic insights in task-trained biology-inspired RNNs with simple tasks (e.g. maintaining persistent representations) is an essential first step to filling this gap, and sets the foundation for scaling these models to challenging tasks that require manipulation of information held in working memory. Examples of complex cognitive functions that interact with working memory include abstract reasoning [21], decision making [22], planning [23], and the execution of behavioral sequences [24].

Motivated by identifying such mechanistic principles, in this study we considered the problem of expanding an RNN to include multicompartment biophysical neurons that have dendrites and active ion channels, and training it to solve a simplistic working memory task that requires maintaining a persistent representation of an external cue (e.g. external input Figure 1A and B). The addition of dendrites with active ion channels is, in our view, the most useful first step to expand biological realism, as dendrites are considered a promising addition for constructing more brain-inspired RNN models, with numerous empirical and computational studies indicating their unique computational benefits for processing task-relevant information from extrinsic sources [25, 26].

**Fig 1.**
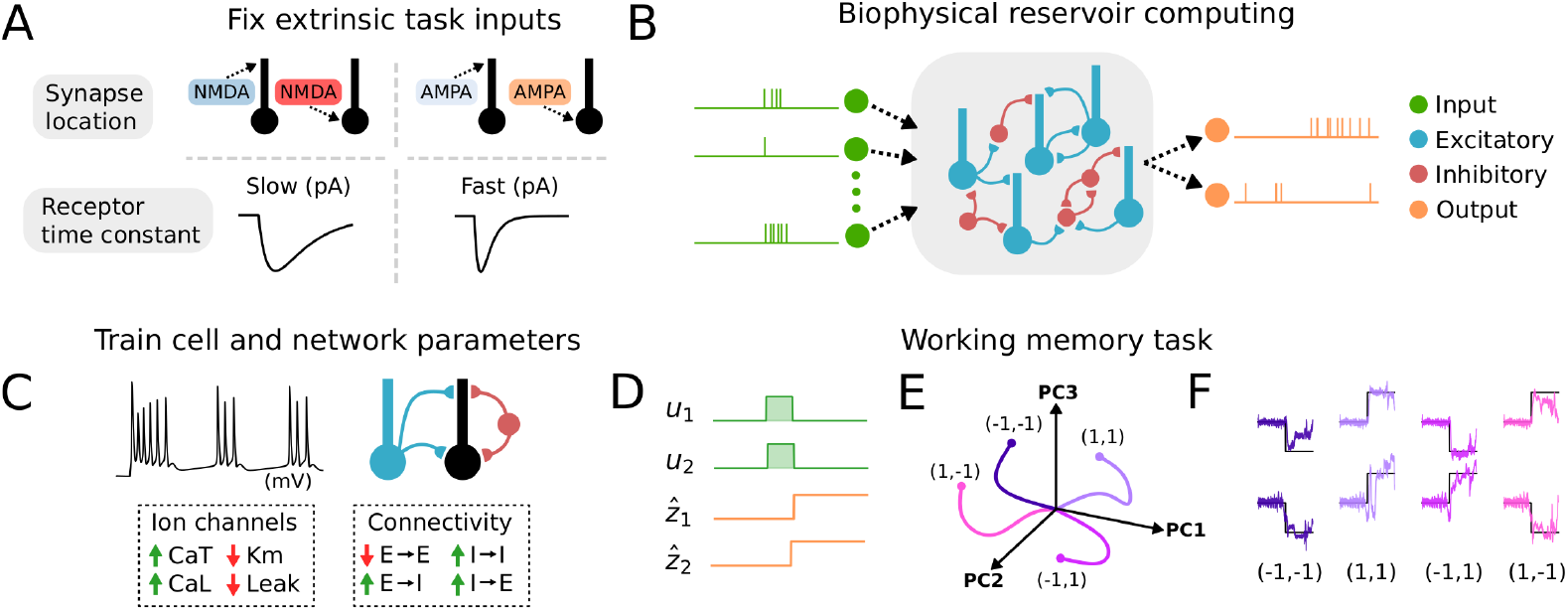
Biophysical reservoir computing. **A:** The goal of this study was to provide mechanistic insights to guide the construction and training of biophysical neural models to solve cognitive tasks. We focused on choices related to providing task-relevant extrinsic inputs to the network. 4 distinct networks were trained to perform a working memory task with fixed extrinsic input properties. These included slow time constant “NMDA-like” inputs at the dendrite or soma (NMDA_dend_ and NMDA_soma_), and fast time constant “AMPA-like” inputs at the dendrite or soma (AMPA_dend_ AMPA_soma_). **B:** Schematic of the biophysical reservoir computer (BRC). The basic building blocks are multicompartment neurons representative of cortical excitatory and inhibitory neurons. Extrinsic inputs are delivered through a population of spiking neurons that are tuned to task-related information. The biophysical reservoir consists of recurrently connected excitatory and inhibitory neurons. Outputs are calculated as a weighted combination of the firing rates of a subset of neurons. **C**: With extrinsic input properties fixed (panel A), the local cell (i.e. ion channel conductance) and network (i.e. synaptic connectivity) parameters of the BRC were optimized for the working memory task. **D:** Cue information for the working memory task (*u*_1_ and *u*_2_) is delivered for a brief 25 ms period. The target output of the network (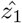 and 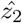) is a 2 dimensional vector that matches the sign of the cue inputs. The goal of the task is to match the target output after the termination of the cue input. **E:** Schematic neural trajectories of a network successfully trained to perform the working memory task. Each trajectory represents a separate trial condition. **F:** Outputs of a network successfully trained to perform the working memory task.

A major contribution of this study is the creation of a novel biophysically-detailed reservoir computing (BRC) framework (Figure 1B), which was specifically developed for deriving mechanistic insights from more biophysical RNN models trained to solve cognitive tasks (i.e. working memory). We directly build upon the biophysical modeling platform Jaxley [27], a powerful software package which applies modern deep learning techniques to the simulation and training of detailed biophysical neural models. Due to the complexity of the BRC network, current optimization methods available in Jaxley (e.g. gradient-based optimization) were insufficient to train the network on the working memory task. Therefore, we developed a novel deep learning based evolutionary algorithm (Figure 2A, see “Neural flow evolution” section of Methods).

**Fig 2.**
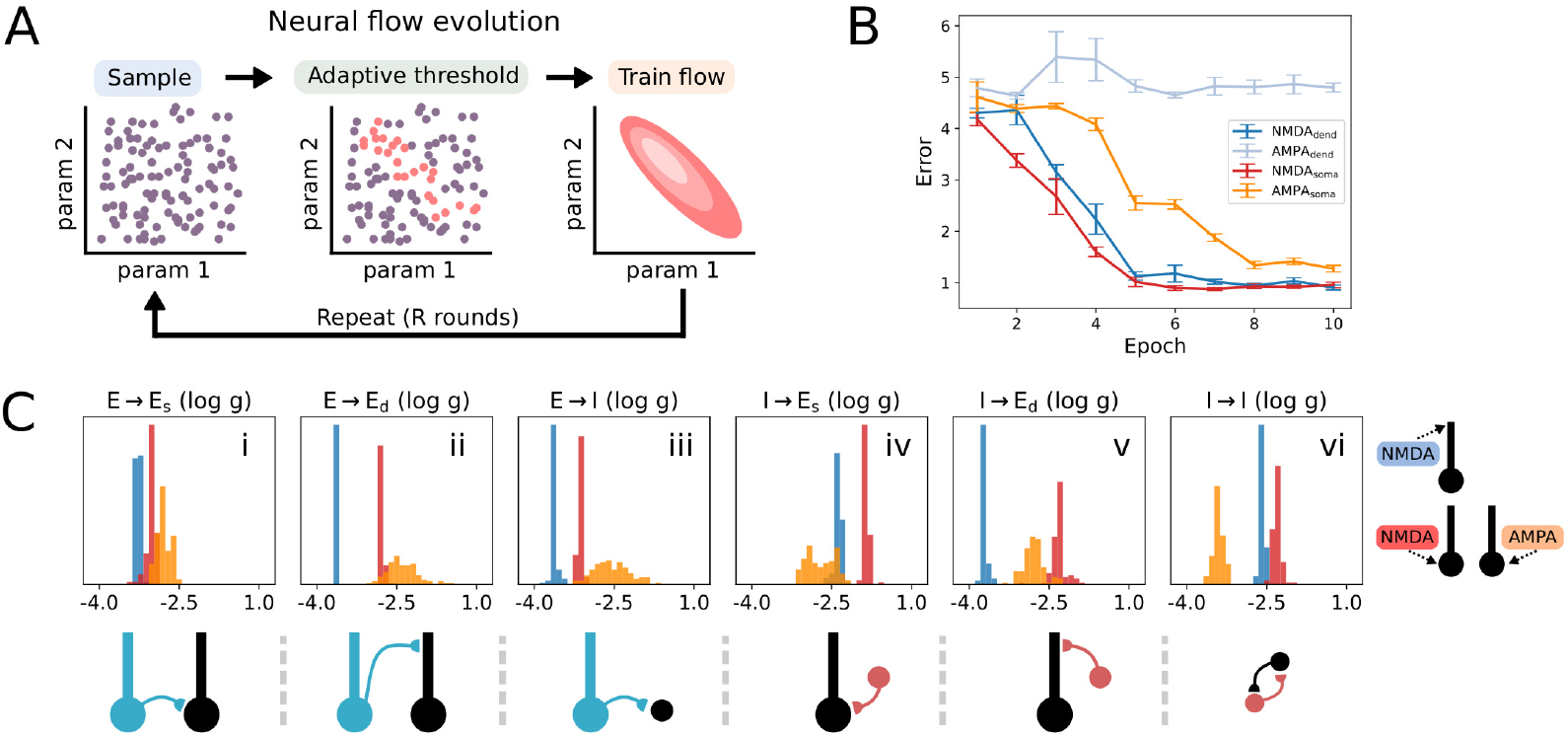
Neural flow evolution. **A:** Schematic of the evolutionary algorithm used to optimize the parameters of the biophysical reservoir computer to perform a working memory task. The initial population of parameters is created by uniformly sampling the parameter space. Each parameter set is simulated and assigned a fitness score for its performance on the working memory task. An adaptive threshold is calculated based on the quantile of the current population’s error. A neural density estimator is trained to approximate the distribution of parameters that remain after thresholding. A new generation is created by sampling from the trained neural density estimator. **B**: Optimization error for NMDA_dend_ (dark blue), AMPA_dend_ (light blue), NMDA_soma_ (red) and AMPA_soma_ (orange) extrinsic input variants. NMDA inputs at either the soma or dendrite rapidly converged to the target error level of 1.0 compared to the AMPA inputs. While AMPA_soma_ eventually converges to the target error level, AMPA_dend_remained at chance level error. **C(i-vi):** Optimized connectivity parameters for the successfully trained network variants NMDA_dend_, NMDA_soma_, AMPA_dend_. Values correspond to synaptic conductance in nanosiemens. Connections shown included E → E_s_ (i), E → E_d_ (ii), E → I (iii), I → E_s_ (iv), I → E_d_ (v), and I → I (vi)

Reservoir computing was intentionally chosen for the purpose of finding network configurations that can perform the task, as opposed to targeted optimization of parameters on a cell-by-cell basis. Reservoir computing refers to a system where input signals are mapped into a high-dimensional space through the activity of a fixed dynamical system (i.e. the reservoir) [28–30]. The training primarily occurs on the linear readout weights, and shared parameters of the reservoir that shape its response characteristics. The rationale is that we are looking to identify computationally useful properties (e.g. connectivity rules, spatial distribution of ion channel conductances, ideal morphologic complexity of dendrites) that can be generalized to any network obeying a limited set of biological constraints. This method is similar to pre-training of deep neural networks, where the goal is to create a generically useful base network that can be fine-tuned to a specific task [31–33].

We hypothesize that expanding task-trained neural models to include dendrites necessitates a careful treatment of 1) the location (e.g. dendrite vs. soma) and synaptic time constants of the extrinsic “cue” input onto the dendrite-equipped neurons, and 2) the intrinsic biophysics (e.g. ionic currents) and local synaptic connectivity of the dendrite-equipped neurons (Figure 1A). These hypotheses are largely motivated by experimental neuroscience indicating the importance of the aforementioned properties for working memory.

With respect to synapse time constants, numerous experimental and computational studies have implicated NMDA receptors as a vital component of working memory in the neocortex [34–37]. The most compelling forms of evidence demonstrate that selective blockade of NMDA receptors in dorsolateral prefrontal cortex impairs persistent firing of cortical neurons [36, 38]. The distinctive property of NMDA receptors is their slow time constants, and magnesium ion block [39]. These properties combine to function as a coincidence detector, where slight depolarization of the postsynaptic neuron is required to remove the magnesium ion block from NMDA receptors. If a presynaptic neuron delivers glutamate to a postsynaptic NMDA receptor, coincident with a slight depolarization of the postsynaptic neuron, then a sustained depolarization occurs due to the slow time constants of NMDA receptors.

With respect to synapse location, NMDA receptors in rodents are observed to be selectively enriched at distal regions of the apical dendrite, and decrease with closer proximity to the soma [40]. These experimental observations suggest that NMDA serves a functional role that is potentially related to the physiology of pyramidal neuron dendrites. Further, numerous studies have emphasized that this spatial localization of NMDA receptors underlies several of the potential computational benefits provided by dendrites [41–45].

While these experimental and computational studies are promising indicators that both synapse location and time constants are critical components of working memory, it is unclear if this conclusion holds in task-trained biophysical models that are not necessarily constrained to match real brain activity. It is important to emphasize that the framework described here was not designed for determining how computations are carried out in real brains. Instead we ask the following question: if you are allowed to recombine cell-level building blocks of biology, which biophysical properties provide task-specific computational benefits that can be transferred to biology-inspired artificial intelligence (AI) systems? While the choice in biophysical properties can be motivated by neuroscience literature, the trained models are not required to mimic realistic brain activity. This allows a disentanglement of the functional role of biophysical properties, from biological constraints imposed by real brains that are unrelated to the cognitive task being considered.

With these considerations in mind, we ask the following questions: Does the synaptic location and time constants of task-relevant extrinsic inputs representing the cue impact the ability of a network to maintain persistent representations? (Figure 1A-B and D-F)? How do these variations in extrinsic inputs impact the learned intrinsic biophysics and local connectivity necessary for persistent representations (Figure 1C)? Answering these questions would provide valuable mechanistic insights for intelligently constructing task-trained biophysical models, and improve training efficiency by providing validated choices for biological details (such as connectivity, receptor time constants, morphological complexity, ion channels, cell heterogeneity, and combinations of these details) thus avoiding choices that are destined to fail due to the unique mechanistic properties of biophysical neural models.

## Methods

### Biophysical reservoir computing

Neuron simulations were carried out with the simulation software Jaxley [27]. The biophysical reservoir consisted of 50 excitatory neurons, and 25 inhibitory neurons (Figure 1A).

Excitatory neurons were modeled such that their main morphologic feature was a large apical dendrite that attenuated synaptic inputs at distal locations (assuming a passive dendrite). This choice was made to focus on networks with neurons that use active ion channels to propagate distal synaptic inputs to the soma. Figure 6A(iv) shows a schematic of the excitatory neuron’s morphology. The morphology of the excitatory neurons was based on an example from [46] (Figure 3 in the corresponding publication) titled “A reduced compartmental model that replicates active dendritic properties”. The excitatory neuron was constructed with 4 compartments total, where each compartment had 4 subcompartments over which the voltage was calculated. The length and diameter of each compartment was as follows: soma (25 ×25 *µ*m), 1st apical segment (100 ×2.5 *µ*m), 2nd apical segment (100 ×1 *µ*m), 3rd apical segment (100×0.5 *µ*m). Inhibitory neurons consisted of a single compartment (with 4 subcompartments) representing the soma, with a length and diameter of (5 ×5 *µ*m).

**Fig 3.**
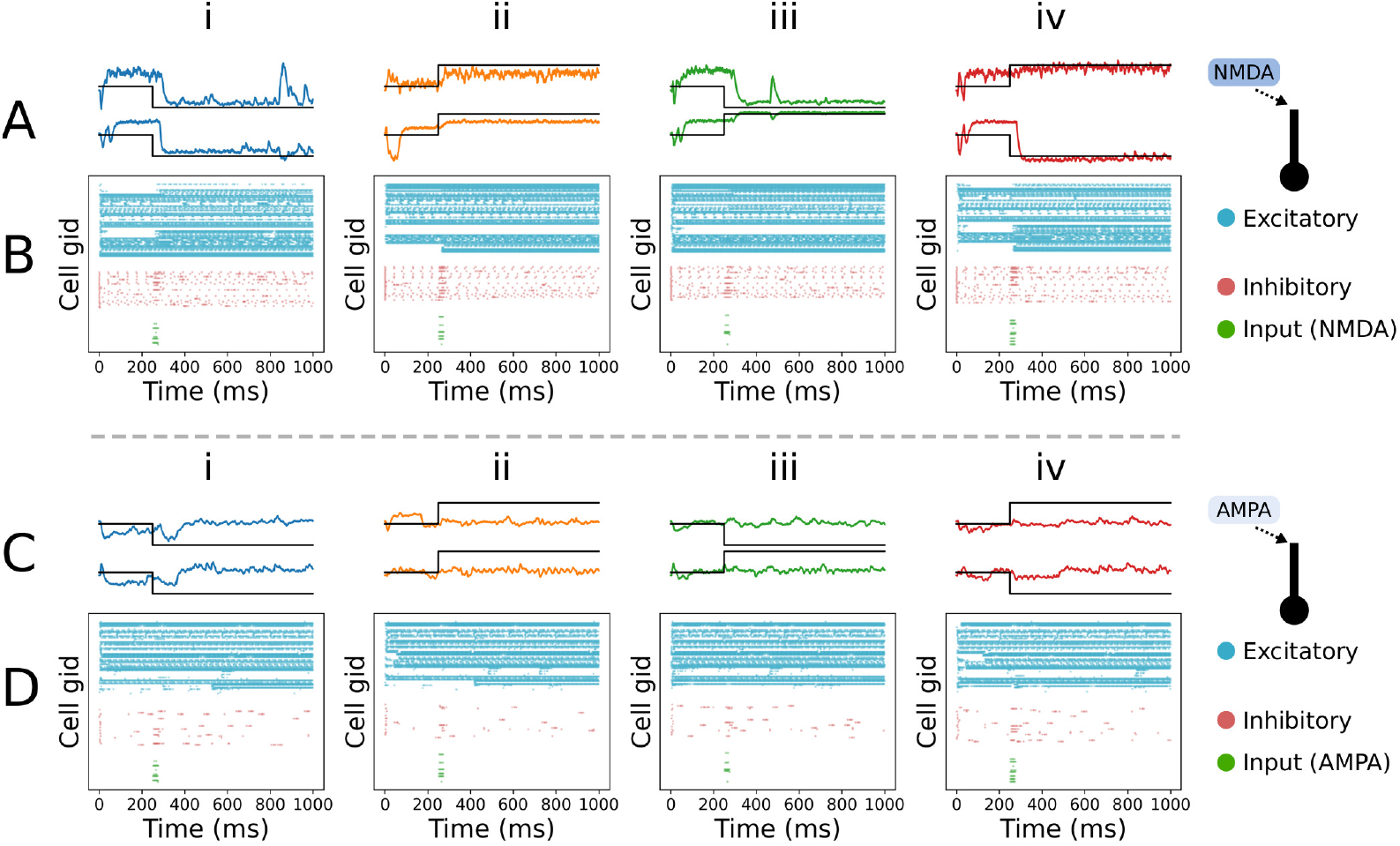
Network-level spiking response to dendritic inputs. **A (i-iv):** Outputs of the trained biophysical reservoir computer with dendritic NMDA inputs (NMDA_dend_). Panels i-iv correspond to the 4 possible types of input stimuli. Model predictions (colored lines) are overlaid on the target output (black lines). As shown, the NMDA_dend_ network was successfully trained to perform the working memory task and maintains outputs in close proximity to the target. **B (i-iv)**: Spike rasters for the simulations underlying the outputs shown in panel A. Input neurons (green) spike briefly during the cue stimulus period (250-275 ms) triggering an immediate change in the spiking of both excitatory (blue) and inhibitory (red) neurons. The change in spiking is unique for all 4 task conditions and persists for the remainder of the 1 second simulation. **C (i-iv)**: Outputs of the biophysical reservoir computer with dendritic AMPA inputs (AMPA_dend_). The optimizer was unsuccessful in training the network with AMPA_dend_(simulations from the best scoring model are shown). Model outputs hover near zero indicating that activity from the network is not distinct across the 4 task conditions, and does not persistently change after the cue inputs. **D (i-iv)**: Spike rasters for the simulations underlying the outputs shown in panel B. Activity of input neurons at 250-275 ms produces no visible change in spiking of excitatory neurons, and a minimal increase in spiking of inhibitory neurons. The change in spiking does not persist after the cue stimulus period, and is not visibly distinct for the 4 task conditions.

Both excitatory and inhibitory neurons were equipped with a diverse set of ion channels available in the base software of Jaxley. The ion channels were based on models described in [47]. These included sodium, potassium, leak, T-type calcium, L-Type calcium, and slow M potassium channels. The maximal conductance 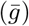 of sodium and potassium channels were fixed to their default values of 0.05 S*/*cm^2^ and 0.005 S*/*cm^2^. The maximal conductance of the remaining ion channels were treated as free parameters to be optimized. The range of values for each parameter are described in Table 1.

**Table 1.** Simulation parameter ranges for neural flow evolution.

| Channel conductance ($\bar{g}$) | | Range | Transform |
| --- | --- | --- | --- |
| E<sub>soma</sub> Leak (S/cm<sup>2</sup>) |  | (-9, -2) | exponential |
| E<sub>soma</sub> Km (S/cm<sup>2</sup>) |  | (-9, -2) | exponential |
| E<sub>soma</sub> CaL (S/cm<sup>2</sup>) |  | (-9, -2) | exponential |
| E<sub>soma</sub> CaT (S/cm<sup>2</sup>) |  | (-9, -2) | exponential |
| E<sub>dend</sub> Leak (S/cm<sup>2</sup>) |  | (-9, -2) | exponential |
| E<sub>dend</sub> Km (S/cm<sup>2</sup>) |  | (-9, -2) | exponential |
| E<sub>dend</sub> CaL (S/cm<sup>2</sup>) |  | (-9, -2) | exponential |
| E<sub>dend</sub> CaT (S/cm<sup>2</sup>) |  | (-9, -2) | exponential |
| I Leak (S/cm<sup>2</sup>) |  | (-9, -2) | exponential |
| I Km (S/cm<sup>2</sup>) |  | (-9, -2) | exponential |
| I CaL (S/cm<sup>2</sup>) |  | (-9, -2) | exponential |
| I CaT (S/cm<sup>2</sup>) |  | (-9, -2) | exponential |
| Connection probability | Range | Transform | Synapse |
| E→E<sub>soma</sub> | (0.0, 0.6) | linear | AMPA |
| E→E<sub>dend</sub> | (0.0, 0.6) | linear | AMPA |
| E→I | (0.0, 0.6) | linear | AMPA |
| I→E<sub>soma</sub> | (0.0, 0.6) | linear | GABAa |
| I→E<sub>dend</sub> | (0.0, 0.6) | linear | GABAa |
| I→I<sub>dend</sub> | (0.0, 0.6) | linear | AMPA |
| Connection strength ($\bar{g}$) | Range | Transform | Synapse |
| E→E<sub>soma</sub> (μS) | (-4, 1) | exponential | AMPA |
| E→E<sub>dend</sub> (μS) | (-4, 1) | exponential | AMPA |
| E→I (μS) | (-4, 1) | exponential | AMPA |
| I→E<sub>soma</sub> (μS) | (-4, 1) | exponential | AMPA |
| I→E<sub>dend</sub> (μS) | (-4, 1) | exponential | AMPA |
| I→I<sub>dend</sub> (μS) | (-4, 1) | exponential | AMPA |
| Cue→E<sub>soma</sub> (μS) | (-4, 1) | exponential | AMPA |
| Cue→E<sub>dend</sub> (μS) | (-4, 1) | exponential | AMPA |
| Cue→E<sub>soma</sub> (μS) | (-4, 1) | exponential | NMDA |
| Cue→E<sub>dend</sub> (μS) | (-4, 1) | exponential | NMDA |
| Cue→I (μS) | (-4, 1) | exponential | AMPA |

The prior support and transform function for the parameters of examples shown in the text. For neural density estimation, all parameters were transformed such that the minimum and maximum of the **Range** column was on the interval $[0, 1]$ .

Our network model included AMPA, NMDA, and GABAa synapses. The synapse models were imported from the Jaxley-mech library (https://github.com/jaxleyverse/jaxley-mech), a companion repository to Jaxley [27]. In general, we used the default parameter values for each synapse, which match the fitted values in [48]. The only exception is for experiments where the time constants of NMDA and AMPA synapses were modified for extrinsic inputs. See the “Extrinsic synaptic receptors” section of Methods for a detailed description. The maximal conductance for each ion channel was set to be equal across all neurons. For excitatory neurons, ion channel maximal conductance parameters were set separately for the soma (1 value for 1 compartment) and the dendrite (1 value for 3 compartments).

Background noise was delivered through AMPA synapses located on the soma of all neurons with spike times generated from a poisson-like process. Each neuron received independent noisy inputs with an average firing rate of 1 Hz and maximum conductance of 1 ×10^*−*4^ nS.

A Watts-Strogatz graph with k=5 neighbors was used to establish the baseline connectivity with a small-world topology [49]. Cortical circuits have been shown to obey small-world connectivity patterns [50], and this unique topology is believed to promote dynamical complexity in neural circuits [51, 52]. Excitatory and inhibitory neurons were recurrently connected through AMPA and GABAa synapses. Synapse locations included the soma and the 3rd apical segment (furthest from soma) for excitatory neurons, and exclusively the soma for inhibitory neurons.

A population of 25 input neurons was used to deliver task-related cue information. Input neurons synapse at 2 possible locations on excitatory neurons (soma and 3rd apical segment). Inhibitory neurons received synaptic input at the soma. Synapse identity to excitatory neurons was either NMDA or AMPA. Input neurons always connected to inhibitory neurons through AMPA synapses. Input neurons were only connected to half of the neurons in the biophysical reservoir. The firing rates of the other half of neurons were used as readouts. This constraint required cue related information to be transmitted through at least one recurrent synapse for successful performance of the task.

The conductance and connection probability of all connections (input and recurrent) was treated as a free parameter to be optimized. The range of values for each parameter are described in Table 1.

All parameters described in Table 1, as well as fixed parameter values, were set to be equal for every neuron belonging to the same class (i.e. excitatory soma, excitatory dendrite, inhibitory, and input neurons). During training with “Neural flow evolution” (see corresponding section below), the parameter updates were applied uniformly across all neurons. The primary sources of stochasticity and heterogeneity in the network were 1) distinct populations of excitatory and inhibitory neurons, 2) noisy background inputs, and 3) probabilistic connectivity.

### Extrinsic synaptic receptors

The kinetics of AMPA and NMDA receptors were modeled according to [48] and [27] as

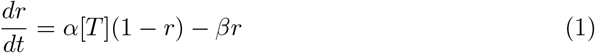

where *r* is the fraction of receptors in the open state, *T* is transmitter concentration, *α* is the forward rate constant, and *β* is the backward rate constant. The AMPA receptor postsynaptic current *I*_*AMP A*_ was calculated as

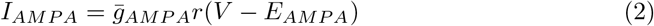

where 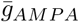 is the maximal conductance, *V* is the postsynaptic membrane potential, and *E*_*AMPA*_ is the reversal potential.

The NMDA receptor postsynaptic current *I*_*NMDA*_ was calculated as

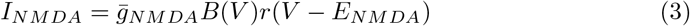

where *B*(*V*) represented a voltage-dependent magnesium block modeled as

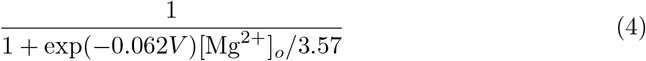

where [Mg^2+^]_*o*_ is the extracellular magnesium concentration. We used the default [Mg^2+^]_*o*_ = 1*mM*.

As described in [48], the forward and backward rate constants for AMPA receptors were set to *α* = 1.1×10^6^*M*^*−*1^*s*^*−*1^ and *β* = 190*s*^*−*1^. The rate constants for NMDA receptors were set to *α* = 7.2 ×10^4^*M*^*−*1^*s*^*−*1^ and *β* = 6.6*s*^*−*1^.

In the final two sections of the results, we investigated what aspect of NMDA receptor properties were important for attractor dynamics. To disentangle the importance of the rate constants from the magnesium block, we ran simulations where *α* and *β* for AMPA and NMDA were swapped. This allowed simulations of a slow “AMPA-like” receptor, and a fast “NMDA-like” receptor.

### Working memory task

A simple working memory task was used to assess the impact of cell-level biophysical properties on network-level dynamics. The task required responding to a cue signal, and maintaining a unique representation of the cue signal until the end of the simulation (Figure 1(D-F)). Cue information was delivered through input neurons activated at 250 ms for a duration of 25 ms. The cue signal consisted of a 2-dimensional vector that remained at zero in the pre-stimulus period, and immediately took on a value *û* ∈ {−1, 1}^2^ in the cue stimulus period for a total of 4 unique task conditions (Figure 1F).

The population of input neurons encoded cue information by mimicking sensory neurons with a gaussian receptive field. Each neuron was assigned a 2-dimensional gaussian distribution with a random mean *µ ∈* [*−*2, 2]^2^ and standard deviation 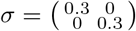. During the stimulus period where cue information was delivered, a 25 ms current injection was applied to the soma of input neurons with an amplitude scaling factor proportional to the gaussian receptive field evaluated at the cue location. The maximum amplitude of the current injection was set to 0.06 nA.

The output *z*(*t*) of the network was defined as a linear readout such that

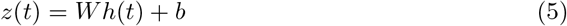

where *h*(*t*) approximates the instantaneous firing rate of excitatory neurons, *W* is a weight matrix of shape *d* ×*n* (*d* = dimensionality of output, *n* = number of excitatory neurons), and b is a bias term. The terms *W* and *b* are learned through linear regression. In this study we used Ridge regression with regularization parameter *α* = 2.0.

Task performance was evaluated with the loss function:

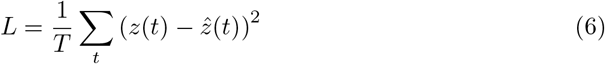

where *z*(*t*) is the network output, 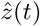 is the target output, *T* is the total number of time steps, and *t* is the current time step. For each set of parameters, the network was simulated with the 4 unique task conditions for 5 trials making a total of 20 simulations per parameter set. The loss used for optimization was calculated as the average loss from leave-one-out cross validation, where 4 of the trials were used to train a linear regression model to predict outputs on the remaining held out trial.

### Neural flow evolution

To optimize parameters of the network model, we took an approach inspired by evolutionary algorithms [53], and combined it with neural density estimation [54] (Figure 2A).

The key steps of an evolutionary algorithm include 1) generate the initial population *θ* ~ *p* (*θ*) through uniform sampling of the prior distribution, 2) evaluate the fitness of the population by simulating parameters *θ*, 3) select the parent population by choosing individuals with high fitness (here we selected parents whose fitness exceeded the 90th percentile of the current generation), and 4) produce offspring by recombining parameters of the parents. For this study, the fitness of each parameter combination (step 2) was evaluated as the negative of the loss function such that the fitness is maximized.

A methodological innovation in this study is the use of neural density estimation [54] to produce offspring from the parent population. Given the parameters *θ* of the parent population, the neural density estimator *q*_*ϕ*_(*θ*) is trained to maximize the average log likelihood:

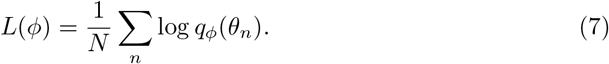

In this study we used a masked autoregressive flow architecture for neural density estimation, and trained the estimator with Adam for 5000 iterations. For each generation, a newly initialized estimator was used. Elitism was implemented in the evolutionary algorithm such that all parameters and simulations were cached in a common dataset during optimization. This allowed parameters from previous generations that exceeded the fitness threshold to be included in subsequent parent populations. To prevent the algorithm from prematurely converging on parameters sets that produced minimal activity, we subtracted a large penalty from the fitness of parameters where the mean absolute value of the output *z* was less than 0.1.

The formal steps of neural flow evolution are as follows:

#### Algorithm 1

Neural flow evolution

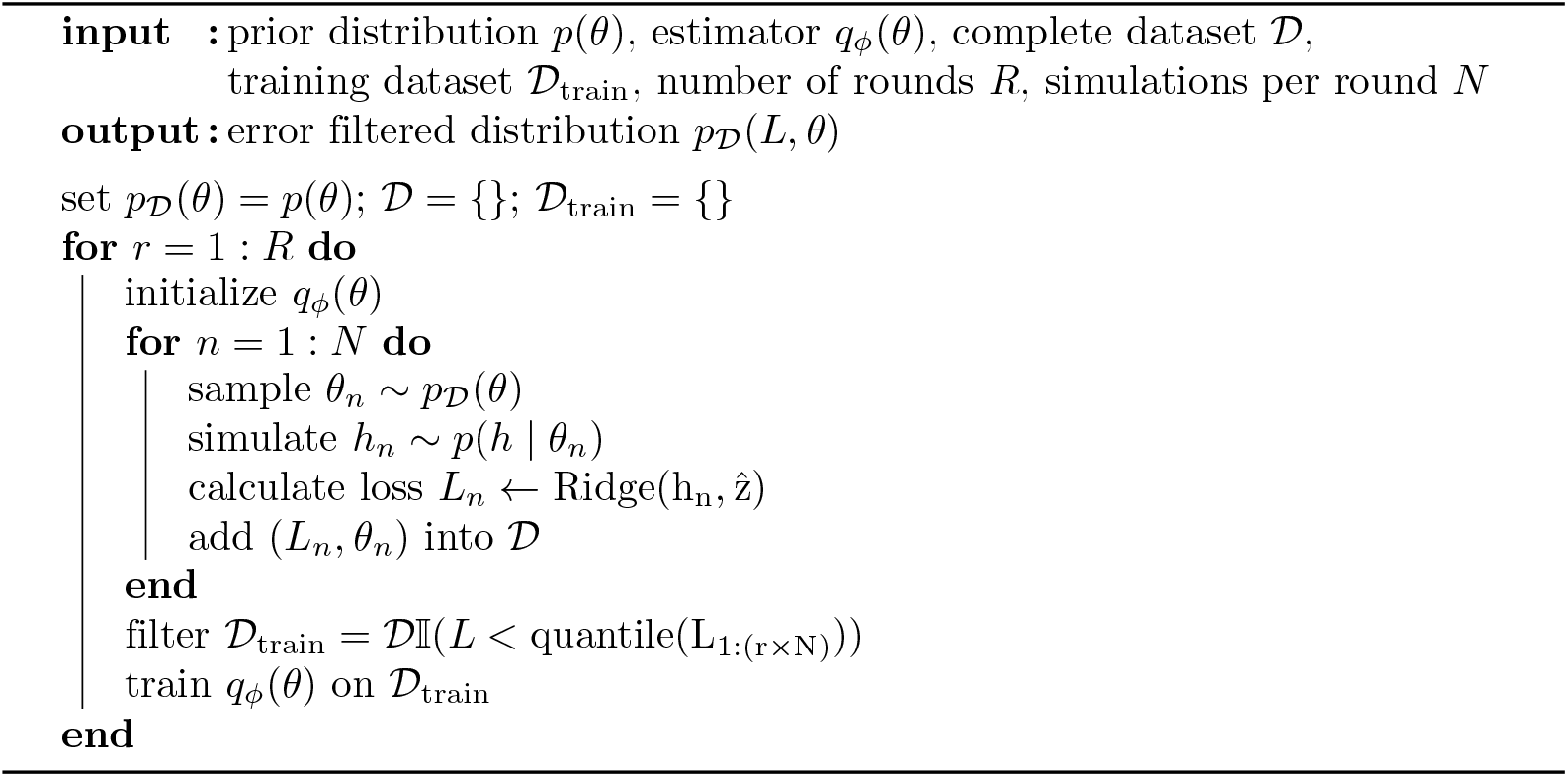

We used a simulation budget of *N* = 100 and trained the networks for *R* = 10 rounds. Each individual “simulation” actually corresponded to 4 separate simulations for each condition in the working memory task. As noted above, loss was estimated using leave-one-out cross validation with 5 repeats. Therefore, 100× 10× 4 × 5 = 20, 000 1 second simulations (differential equation solver time step *dt* = 0.025 ms) were run for the optimization of each extrinsic input variant. Simulations were run on GPU nodes available on Brown University’s high-performance computing cluster (Oscar). Optimization of an individual extrinsic input variant on a single GPU took approximately 15 hours using the optimization parameters described above.

### Overlap coefficient

To compare the optimized parameter distributions learned by the estimator *q*_*ϕ*_(*θ*), we used the distribution overlap coefficient (OVL) [55, 56] which is calculated as:

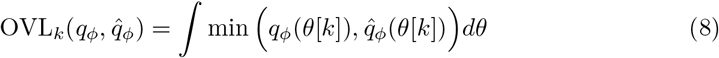

where *q*_*ϕ*_(*θ*[*k*]) and 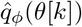 are two different distributions we seek to compare on the marginal distribution for the *k*-th parameter. To calculate the OVL_*k*_ numerically we used an evenly spaced grid of size 30^*d*^ over *d* ∈ ℕ_+_ parameters. The OVL varies on a scale of [0,1] such that 1 indicates complete overlap, and 0 indicates no overlap.

## Results

### BRC reveals that NMDA receptors are more efficient than AMPA receptors in solving working memory tasks

We applied biophysical reservoir computing (BRC) to study how synapse location and receptor time constants of extrinsic inputs impacted a network’s ability to solve a simplified working memory task. We tested 4 extrinsic input variants using the biophysical reservoir computing (BRC) framework, where the properties of task-relevant extrinsic inputs were fixed (Figure 1A), and intrinsic biophysics and connectivity were freely optimized (Figure 1C). In each network variant, the synapse location of extrinsic inputs to excitatory neurons was either the soma or the 3rd apical segment (furthest from the soma), and the synapse time constants were either slow (“NMDA-like” NMDA_dend_ and NMDA_soma_) or fast (“AMPA-like” AMPA_dend_ and AMPA_soma_).

We trained each of the 4 extrinsic input variants on the working memory task (see “Working memory task” section of Methods), and examined the resulting population errors. Figure 2B plots the average population error for each extrinsic input variant over 10 epochs. As shown, AMPA_dend_ was the only network that failed to reach the target error threshold of 1.0, and both of the NMDA conditions reached the target threshold faster than the AMPA_soma_ condition. More specifically, both NMDA_dend_ and NMDA_soma_ converged to the target error within 5 epochs, while AMPA_soma_ converged to the error threshold within 8 epochs. Further, for networks with inputs on the dendrites, NMDA receptors are required for learning that task, as networks with dendritic AMPA receptors were unable to learn the task. For task-related inputs on the soma, either NMDA or AMPA receptors were sufficient for performing the working memory task, but NMDA outperformed the AMPA receptors.

To gain mechanistic insight into how the network learned the task, we also examined the resulting local synaptic connectivities after training in each extrinsic input variant that successfully learned the tasks (NMDA_dend_, NMDA_soma_, AMPA_soma_) to assess if and how the connectivities needed to be set up to solve the working memory task. By plotting histograms of the learned synaptic weights of local connections, it is possible to identify distinct network configurations (local synaptic connectivity patterns) for each extrinsic input variant when their corresponding histograms are non-overlapping. Differences between parameter distributions were quantified using the overlap coefficient (OVL, see “Overlap coefficient” section of Methods), where an OVL of 0.0 indicates a large separation between distributions, and an OVL of 1.0 indicates complete overlap.Figure 2C plots the distribution of learned synaptic strengths for all local recurrent connections in the successfully trained extrinsic input variants (NMDA_dend_ (dark blue), NMDA_soma_ (red), and AMPA_soma_ (orange)).

As shown, the extrinsic input variants produced highly distinct network configurations. Synaptic targets of connections described below are abbreviated with E_s_ (excitatory soma), E_d_ (excitatory dendrite), and I (inhibitory). Across the learned local connection weights, every variant exhibited a high separation from the other two variants for at least one synapse. The only connection which exhibited low separation for all 3 variants was E→ E_s_ (Figure 2C(i)).

We were specifically interested in understanding if learned network configurations were different with respect to extrinsic input location (soma vs dendrite), and time constants (AMPA vs NMDA). Starting with extrinsic input location, we can compare the dendritic input variant NMDA_dend_, to the somatic input variants NMDA_soma_ and AMPA_soma_. For the connections E → E_d_ (Figure 2D(ii)) and I → E_d_ (Figure 2D(v)),NMDA_dend_ exhibited much lower weights compared to either NMDA_soma_ or AMPA_soma_ such that the histograms had no overlap (OVL=0.0 for NMDA_dend_ vs NMDA_soma_ or AMPA_soma_). It is interesting to observe that these strongly decreased connections occurred at the dendrite, suggesting that reducing the strength of local dendritic inputs is necessary when delivering task-relevant extrinsic inputs to the dendrite.

For comparing network configurations with respect to extrinsic input time constants (NMDA vs AMPA), we observe that networks trained with NMDA_dend_ and NMDA_soma_ produced substantially lower synaptic strengths for all excitatory connections compared to AMPA_soma_ (Figure 2C(i-iii)). We additionally observe that nearly all inhibitory connections for NMDA_dend_ and NMDA_soma_ are higher than AMPA_soma_ (Figure 2C(iv-vi)), with the exception of I → E_d_ (Figure 2C(v)) where NMDA_dend_ has substantially lower weights compared to the other extrinsic input variants as described above, suggesting this connection is less useful for learning. We additionally note a large separation between both NMDA variants and AMPA_soma_ for the I → I connections (Figure 2C(vi)). In fact, this was the only connection in which AMPA_soma_ had an OVL of 0.0 with both of the NMDA network variants. Previous studies have implicated I → I connections as being important for working memory tasks [57], potentially suggesting a relationship between NMDA transmission and inhibitory connectivity for the ability to perform working memory tasks. However, we caution overinterpretation of the optimized parameter values in the context of neuroscience. Our goal was not to construct a model that directly explains how real brains function, but instead identify computational principles inspired by the brain. Nevertheless, the BRC framework could be used in follow up studies to characterize I→ I connections when they are fixed to specific values in the same way that extrinsic inputs are treated in this study.

In summary, training the BRC with neural flow evolution revealed that the synapse location and receptor time constants of task-relevant extrinsic inputs have a strong impact on the learned network configurations necessary for working memory. Further, networks with AMPA_dend_ (i.e. fast time constants delivered at the dendrite), were completely unsuccessful in learning the working memory task, despite the intrinsic biophysics and connectivity being freely optimized. While there are potential computational benefits of dendritic information processing, we demonstrate a constraint on task-related dendritic function, such that NMDA receptors are required to transmit cue information through dendrites for working memory, and overall NMDA seems to be more efficient than AMPA for working memory tasks.

For the remaining sections, we will attempt to understand how these differences emerge with analysis techniques traditionally employed by systems neuroscientists, along with a description of their biological significance. We note that while we adopt tools from experimental neuroscience, our ability to make inferences from simulated neural activity are much stronger than in vivo activity, as we have direct access to any measurable property of all neurons simulated.

### Successfully trained networks produce distinct and persistent changes in network-level spiking activity

In the previous sections, we examined the impact of variants in extrinsic cue input location and time constant on training in the working memory task by inspecting errors in task performance, and the learned connection strengths of the local network. We now move to analyze the cell spiking activity during the working memory task. Spiking activity is essential to characterize when attempting to understand how a biophysical neural network solves a computational task, as spikes indicate the exact moment at which communication between cells is occurring. Similar to experimental neuroscience, by inspecting ongoing spiking activity, we will uncover differences in spiking mechanisms underlying the ability to maintain persistent representations in the BRC network, and identify which aspects of spiking activity indicate good performance.

Empirical studies have shown that working memory processes in frontal cortical networks require both 1) changes in spiking activity that are distinct for different sensory-mediated extrinsic inputs, and 2) a persistent change in spiking activity that lasts after the extrinsic input ends [34, 36]. As such, we were interested in examining how differences in extrinsic inputs (and their optimized intrinsic parameters) impact spiking dynamics to mediate task performance, and if the principles that emerge were consistent with the prior empirical studies. We specifically asked if synapse location and receptor time constants of extrinsic inputs influence network-level spiking response before and after the stimulus, and if the optimized intrinsic parameters for each extrinsic input lead to differences in overall spiking activity throughout the simulation.

Figure 3 and Figure 4 plot spiking activity in each neuron (panels B and D) and the corresponding network output (panels A and C) for a single trial across all task conditions (panels i-iv). Network outputs were produced by a learned set of linear weights (see “Working memory task” set of Methods, outputs for a held out validation set are shown). The simulations were generated from the best performing parameter set identified during optimization. In Figure 5A and B below, we provide a more detailed view by plotting the spiking response of individual neurons across all task conditions.

**Fig 4.**
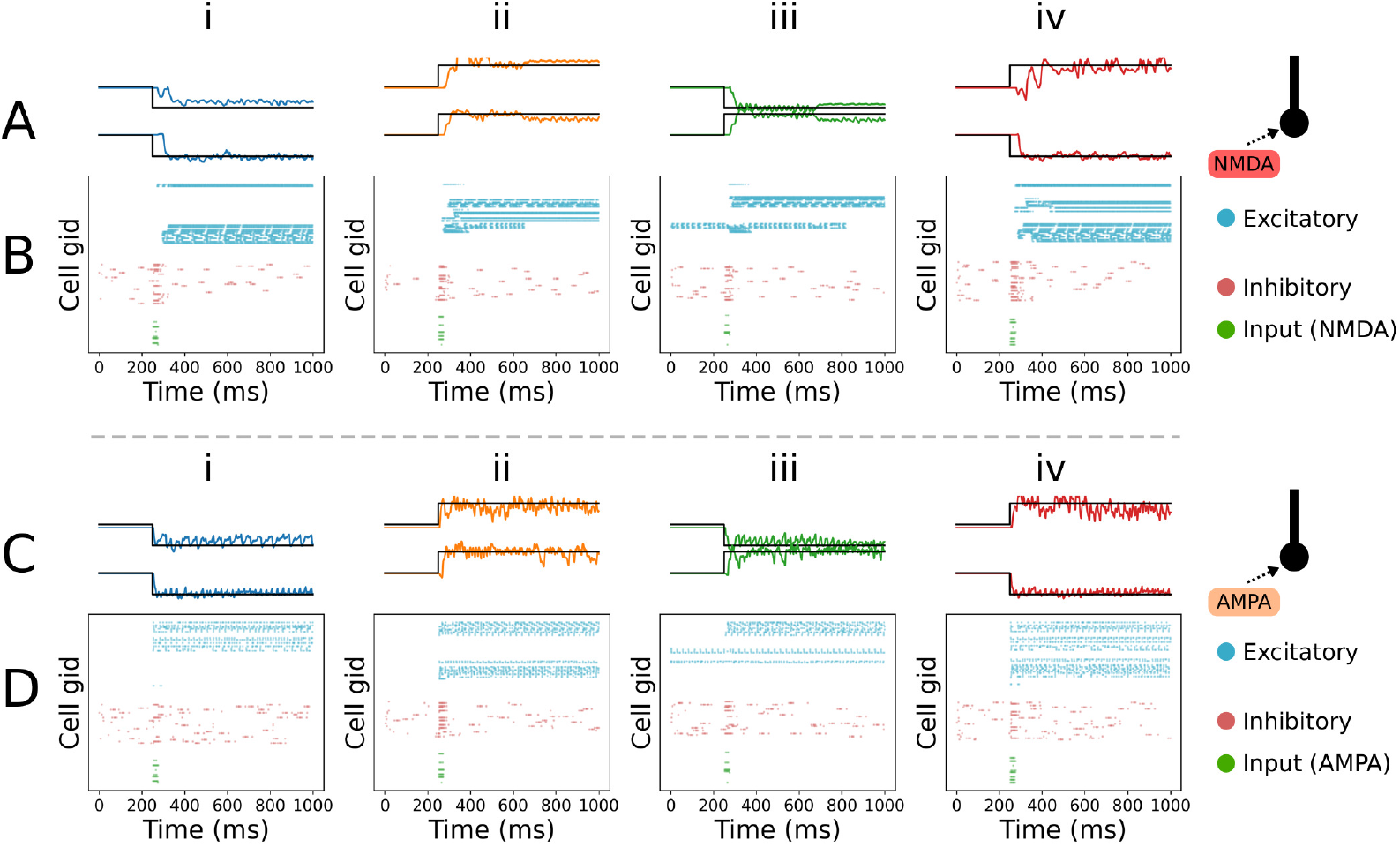
Network-level spiking response to somatic inputs. Figure panels follow the same structure as Figure 3. **A-B**: Model outputs (A) and spike rasters (B) are shown for the network trained with NMDA_soma_ inputs. Across all task conditions (i-iv), the network produces a robust response to the cue input, with distinct clusters of excitatory neurons that maintain persistent firing to the end of the simulation. **C-D**: Model outputs (C) and spike rasters (D) are shown for the network trained with AMPA_soma_ inputs. Similar to NMDA_soma_ (panels A-B), the network exhibits clustered excitatory neuron activity that is distinct for each task condition. bars), this activation has no sustained impact in the post-stimulus period.

**Fig 5.**
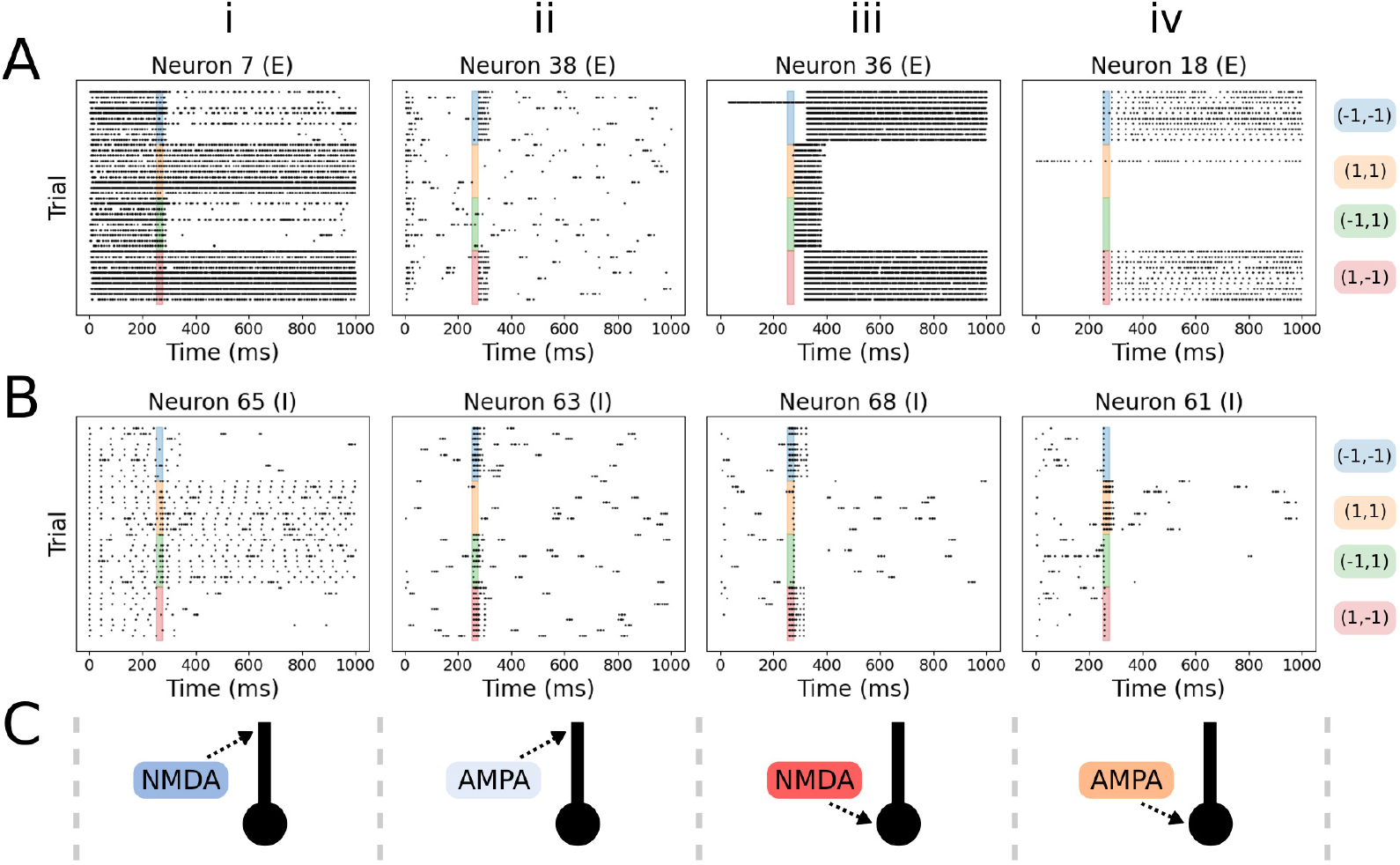
Dendritic NMDA input triggers sustained persistent firing. **A**: Raster plots for individual excitatory neurons during the working memory task. Colored bars indicate the task condition (10 trials per condition). As shown, the successfully trained networks with NMDA_dend_ (i), NMDA_soma_ (iii), and AMPA_soma_ (iv) all exhibit condition-dependent spiking activity. In the network with AMPA_soma_ (ii), which was not successfully trained to perform in the working memory task, the exemplar excitatory neuron exhibits a condition-dependent increase in firing rate when the cue input is active (increased for blue/red, no increase for orange/green). However, the change in activity does not persist past the cue stimulation period (250-275 ms). **B:** Raster plots for individual inhibitory neurons. As shown, inhibitory neurons in successfully trained networks (i, iii, and iv) exhibit condition-specific changes in activity that are sustained after the cue stimulation period. However, in the unsuccessfully trained network with AMPA_soma_ (ii), the inhibitory neuron only shows spiking in response to background noise that does not depend on the task condition. **C**: Schematic of the cue synapse identity and location for each corresponding column.

The networks that were successfully trained to perform the working memory task (NMDA_dend_ in Figure 3A, NMDA_soma_ and AMPA_soma_ in Figure 4A and C) all show network outputs (colored lines) that closely match the target signal (black lines). In contrast, AMPA_dend_, which was not successfully trained, produces outputs with small fluctuations that are uncorrelated with the target signal (Figure 3C).

Inspection of the spike raster plots reveals why AMPA_dend_ exhibits poor performance on the task (Figure 3D). Despite the presence of persistent spiking activity at baseline in excitatory (blue) neurons, activity of the input neurons (green) during the cue stimulus period (250-275 ms) fails to produce any time-locked change that lasts to the end of the 1 second simulation. A small increase in inhibitory (red) neuron spiking aligned to the cue stimulus is observed, but this change is transient as inhibitory neurons primarily exhibit stochastic spiking due to the noisy background input. We hypothesize that strong and synchronous inhibition is key to resetting the spiking activity in the excitatory neurons, permitting the formation of newly activated clusters that encode the memory. This strong and synchronous inhibition does not occur in this case of AMPA_soma_ (Figure 3D), in contrast to NMDA_dend_ and AMPA_soma_ which both exhibit a strong recruitment of inhibitory neuron spiking that is time-locked to the cue input (Figure 3B and Figure 4B and C).

We also observe that the excitatory neurons in AMPA_dend_ seem to have no clear change in spiking aligned to the cue stimulus (Figure 3D). This is in contrast to the NMDA_dend_ network variant where in addition to a cue-aligned change in both excitatory and inhibitory neuron spiking, there is relatively higher inhibitory neuron spiking at baseline (Figure 3B red dots prior to 250 ms). Figure 5B(ii) shows this effect with a spike raster for a single inhibitory neuron, and demonstrates that despite the neuron being selectively activated by input neurons during different trial conditions (colored bars), this activation has no sustained impact in the post-stimulus period.

In the successfully trained networks (NMDA_dend_, NMDA_soma_, and AMPA_soma_), it is clear that the spike rasters demonstrate persistent spiking with distinct responses for each of the trial conditions that lasts to the end of the simulation (Figure 3B and Figure 4B and D). While the spike rasters for NMDA_dend_ exhibits elevated activity across all neurons in the network (Figure 3B), the cue inputs are successfully able to produce lasting changes in both excitatory and inhibitory neuron activity, with distinct clusters entering an “on” state in the post-stimulus period. In contrast, spike rasters for NMDA_soma_ and AMPA_soma_ exhibit less activity across the entire network, in favor of smaller clusters of excitatory neurons that are strongly activated in response to the cue inputs (Figure 4B and D). Since all extrinsic input variants are optimized with the same parameter ranges for local connectivity and biophysics, the elevation in baseline activity for NMDA_dend_ and AMPA_dend_ suggests that the cell and network adaptations necessary for propagating inputs from the dendrites to the soma make the neurons more sensitive to the noisy background inputs, leading to higher spiking at baseline. For example, the changes in biophysics necessary for dendritic spikes may make the neuron more excitable, whereas the biophysics of NMDA_soma_ and AMPA_soma_ are able to suppress responses to noisy inputs, and still produce persistent spiking in response to strong and synchronous cue inputs.

Figure 5 highlights the diversity of single neuron responses across the trained networks. While the spike raster of NMDA_dend_ in Figure 3B clearly demonstrates cue neurons triggering persistent spiking, Figure 5A(i) and B(i) reveals that persistent firing in excitatory and inhibitory neurons can be turned off by the cue inputs. This is an important observation that is similar to modeling work in [58] which also describes a biophysical mechanism in which excitatory inputs can terminate persistent firing. One of the open questions in working memory research is how/why the prefrontal cortex maintains the same overall level of spiking activity when comparing rest to working memory periods [34]. The ability of NMDA_dend_ to turn off persistent spiking in specific neurons offers a possible explanation, such that different sets of neurons are switched on/off under different task conditions, but the overall number of “on” neurons remains the same.

Across the successfully trained networks, inhibitory neurons exhibit low firing rates relative to excitatory neurons (Figure 3B and Figure 4B and D, red dots), except for the initial burst of spikes time-locked to the cue input. In these network-level raster plots, it is not visually apparent if the inhibitory neurons exhibit task condition specific changes in firing activity. However, visualizing spike rasters at the single neuron level demonstrates that the inhibitory neurons still do exhibit lasting changes in spiking that are specific to the task condition (Figure 5B(i, ii, and iv)).

### Both intrinsic bursting and network connections contribute to working memory performance

As described previously, one reasonable hypothesis on why NMDA receptors are useful for working memory is that they are able to trigger persistent spiking activity in single neurons that is maintained during the memory period [34, 38]. Due to the slow time constants of NMDA receptors, extrinsic inputs equipped with NMDA are capable of producing sustained depolarization in the target neurons, with a burst of spiking activity even in the absence of inputs from the local network. Therefore, a simple solution to the working memory task would be for cue evoked extrinsic inputs to act like an “on” or “off” switch, where the input triggers a long burst of spiking in target neurons. If cell ion channels are set up such that the burst does not terminate, then this would be a situation where single neuron biophysics (and not network-level interactions) are sufficient for persistent spiking activity underlying working memory. Further, even if the target neuron is unable to maintain persistent spiking independently, a long burst of spikes could more effectively excite neurons in the local network, and reciprocally excite target neurons to produce persistent spiking.

To examine this possibility further, we assessed if the variants in task-relevant extrinsic inputs (and their optimized intrinsic biophysics) produced distinct burst responses in isolated (**disconnected**) excitatory neurons (Figure 6A). We further asked if the burst responses of isolated neurons were predictive of task performance (i.e. task errors quantified in Figure 2B).

**Fig 6.**
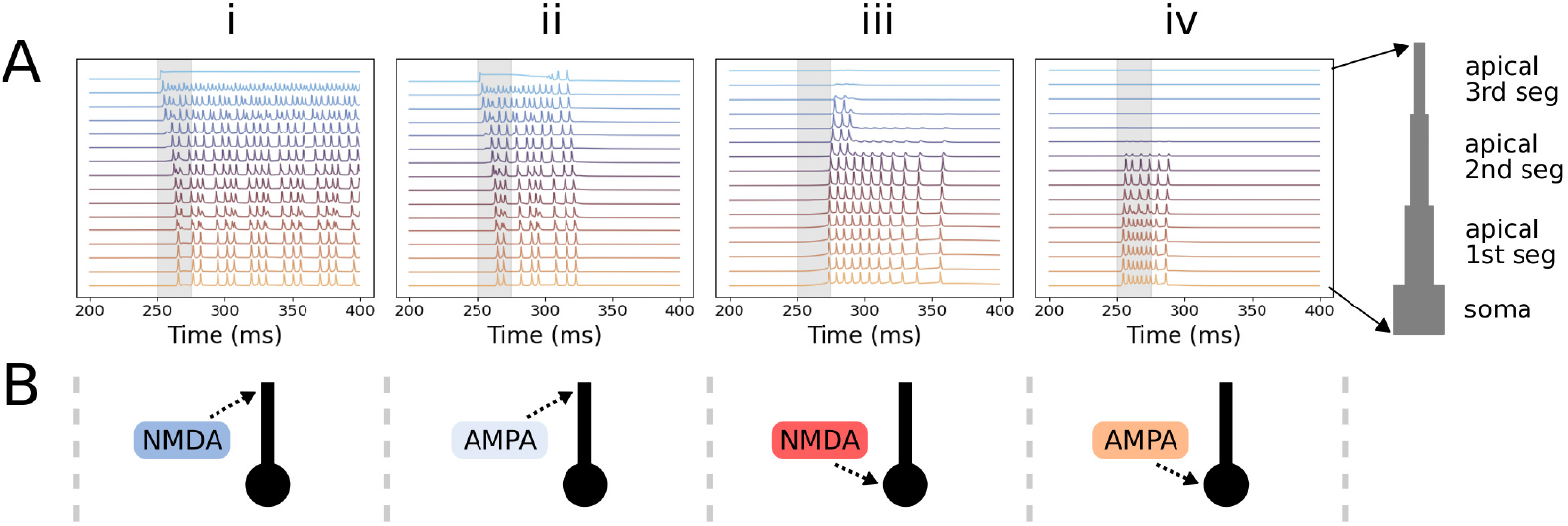
Extrinsic input variants generate diverse lengths of spike bursts. **A**: Membrane potentials of excitatory neurons were recorded in every compartment. Panels indicate the extrinsic input variant simulated, including: NMDA_dend_ (i), AMPA_dend_ (ii), NMDA_soma_ (iii), AMPA_soma_ (iv). The simulations were run with all recurrent connections removed, such that the only synaptic input comes from cue neurons. The 25 ms cue stimulus period is plotted in grey. Panel iv includes a schematic which maps membrane potential traces to each compartment. As shown, all extrinsic input variants initiate a burst of spikes that starts in the cue period (grey bar), and continues for varying amounts of time beyond the end of the cue period. Inputs to the dendrite generate bursts of spikes that propagate via dendritic spikes down to the soma (panel i and ii). Inputs to the soma also generate bursts of spikes, but they fail to propagate to the top of the dendrite (iii and iv). **B**: Schematic of the extrinsic input variant for each corresponding column.

Here, we removed the local connectivity to distinguish how much of the persistent spiking observed in the connected network in Figure 5 was due to inputs from the local network vs a self-generated burst of spiking due to the cell’s own biophysics. During these simulations, the membrane potential was recorded in every compartment of excitatory neurons. We defined a burst as spiking activity that continues autonomously after the cessation of cue inputs. As shown in Figure 6A, the NMDA_dend_ (panel i) and NMDA_soma_ (panel iii) excitatory neurons produce robust bursts of spiking lasting *>* 100 ms, which is substantially longer than the 25 ms cue stimulus period (grey bar). In NMDA_dend_ (panel i) the spiking is initiated in the dendrite by a sustained depolarization of the 3rd apical segment (top-most voltage trace), with dendritic spikes propagating down the apical dendrite to trigger somatic spiking (bottom-most trace).NMDA_soma_ (panel iii) exhibits back propagating action potentials originating from the soma, but they do not completely ascend to the top of the apical dendrite.

Despite the fast time constants of AMPA receptors, extrinsic AMPA inputs to the dendrite and soma were also able to generate single neuron spike bursts (Figure 6A(ii and iv)). In contrast to the burst responses triggered by NMDA inputs, AMPA_dend_(panel ii) and AMPA_soma_ (panel iv) exhibit substantially shorter bursts of ~ 40 ms and ~ 60 ms respectively. Interestingly, while AMPA_soma_ produced the shortest burst length of ~ 40 ms (Figure 6A(iv)), it is clear that when simulated in a connected network, the neuron can produce sustained firing in response to cue inputs (Figure 5A(iv)). This suggests that the ability of AMPA_soma_ to maintain unique representations of cue information highly depends on local recurrent connections in the network, and cannot be solely explained by the bursting properties of individual neurons. Further, despite AMPA_dend_having the same recurrent connections and a similar burst response to cue inputs (Figure 6A(ii)), condition-specific persistent firing does not arise.

Overall, these results demonstrate that the variants of task-relevant extrinsic inputs do indeed produce different bursting activity in isolated neurons, with NMDA receptors producing longer bursts when synapses are located either at the dendrite or soma. Further, despite dendritic AMPA receptors being capable of generating bursting activity that persists beyond the cue stimulus period, the AMPA_dend_network was unable to learn the working memory task. These results suggest that single-neuron firing properties (i.e. bursting) are not entirely predictive of persistent-firing activity that maintain task-relevant information. In the context of scaling biophysical network models to solve more difficult cognitive tasks, these results indicate that researchers should not necessarily begin by optimizing single neuron biophysics for desired response patterns, but instead should optimize cell and network level properties jointly.

### Dendritic AMPA receptors are insufficient for establishing stable attractor dynamics

An increasingly popular way to analyze the dynamics of a simulated neural system is to study how the computations correspond to trajectories that evolve upon a low-dimensional neural manifold [59, 60]. One common dynamical systems-based theory that seeks to explain working memory posits that neural networks flexibly create fixed point attractors which pull neural trajectories along manifolds towards their basin of attraction [61]. Memories correspond to neural trajectories that remain in the basin of attraction for a sustained period of time. Figure 7B provides a schematic relevant to the working memory task in this study, and shows how basins of attraction corresponding to each task condition occupy distinct regions of the state space. Under this framework, the ability of a network to perform a working memory task relates to its ability to establish temporally stable basins of attraction that are separated in the state space.

**Fig 7.**
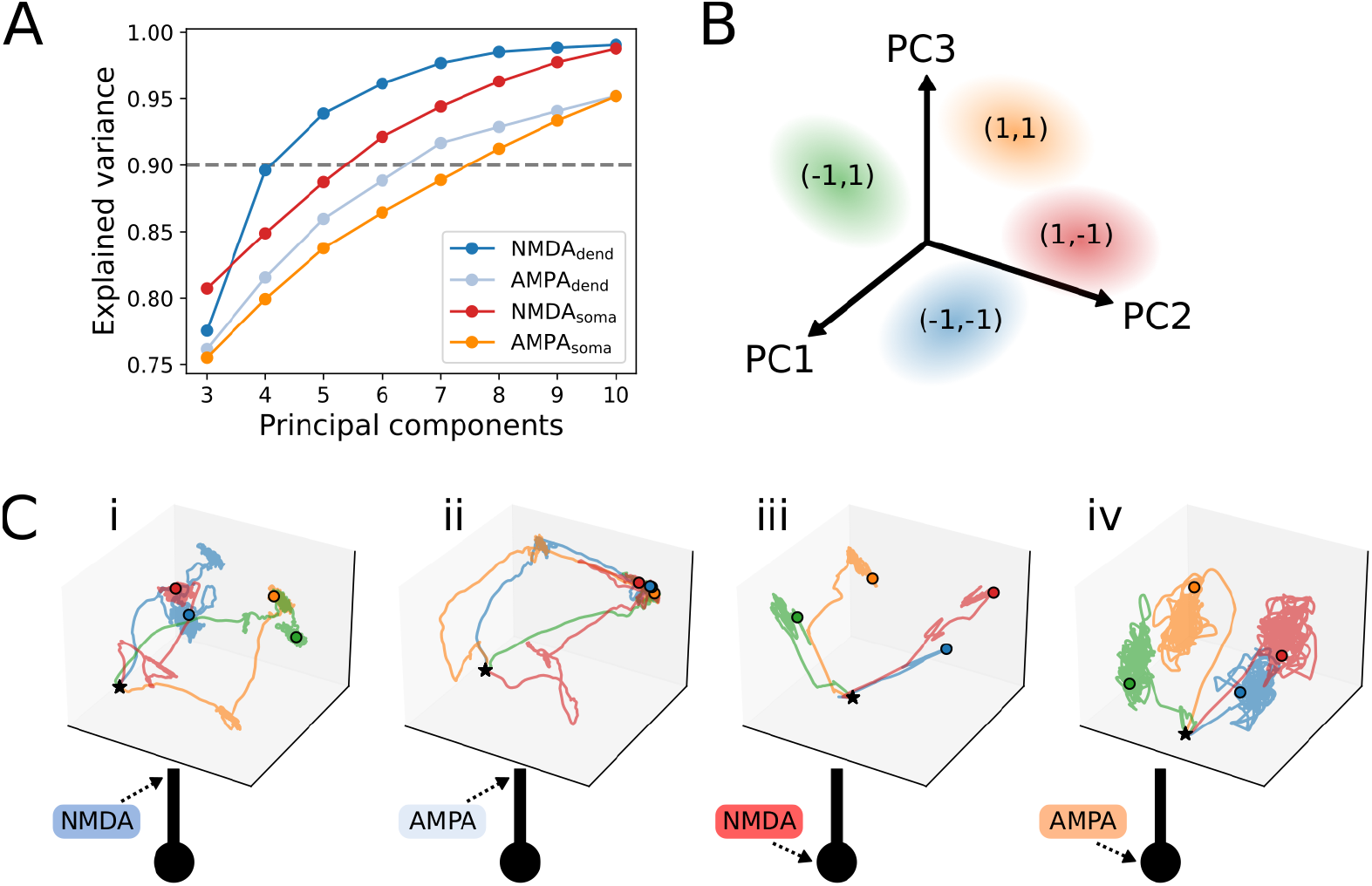
Synaptic receptor identity and location alters attractor dynamics. Network activity was analyzed with principal component analysis (PCA) to assess the impact of the different types of cue inputs on network dynamics during the working memory task. PCA was only applied to the firing rates of excitatory neurons. **A:** Cumulative variance explained for simulations with dendritic NMDA_dend_ and AMPA_dend_, and somatic NMDA_soma_ and AMPA_soma_ cue inputs. Networks with NMDA exhibit lower dimensionality compared to networks with AMPA inputs, as NMDA_dend_ and NMDA_soma_ reach 90% explained variance (dashed grey line) with fewer components than AMPA_dend_ and AMPA_soma_. **B**: Schematic of PCA figures shown in panel C. Colored ovals represent basins of attraction which pull neural trajectories into different regions of the state space depending on the task condition. **C**: Neural trajectories for PC1-PC3 are shown for the 4 extrinsic input variants. Each trajectory corresponds to the entire 1 second simulation. The beginning of the simulation is marked by a black star, and the end of the simulation is marked by a colored dot. The color of the trajectory corresponds to the different trial conditions.

We first asked if the location and time constants of extrinsic synapses impacted the complexity (i.e. dimensionality) of network-level neural dynamics by quantifying its dimensionality with principal component analysis (PCA). In the context of this study, dimensionality is a measure of describing the complexity of the manifold that neural trajectories traverse during the working memory task. Numerous studies have suggested that dimensionality reflects distinct computational processes [62–64]. Given the striking differences in task performance and network-level spiking across extrinsic input variants (Figure 3 and Figure 4), we reasoned that the dimensionality may explain these differences with respect to the complexity of their associated neural manifolds.

Explained variance of the principal components (PCs) was used to estimate the dimensionality of neural activity. Figure 7A plots the variance explained for all 4 variants of extrinsic inputs. Despite the failure of AMPA_dend_ to perform the working memory task (Figure 2B and Figure 3C), the variance explained for each PC is not a large outlier, and instead fits between the curves of NMDA_soma_ and AMPA_soma_. There is a general trend for AMPA inputs exhibiting slightly higher dimensionality, with AMPA_dend_ and AMPA_soma_ reaching 90% variance explained with 7 and 8 PCs respectively, in contrast to NMDA_dend_ and NMDA_soma_ with 4 and 6 PCs respectively. In summary we find that PCA-estimated dimensionality does not directly relate to performance on the working memory task, but instead seems to reflect activity produced by networks trained with either NMDA or AMPA for extrinsic inputs.

Previous researchers have shown that in vivo, the dimensionality of neural activity is strongly modulated by the behavioral task considered, and consequently the extrinsic inputs driving activity in the recorded neural population [64, 65]. Here, we are analyzing the dimensionality of neural activity driven by 4 distinct cue inputs (representing a 2-dimensional target signal) in addition to noise. While the minimal dimensionality of neural activity needed to solve the task is 2 dimensions (or even 1 dimension if the 4 basins of attraction are mapped to different positions on a single axis), it is clear across all extrinsic input variants that the neural activity is explained by substantially more than 2 PCs (the lowest dimensionality was NMDA_dend_ with 4 PCs explaining 90% of the variance), potentially suggesting that more complex activity patterns are necessary to solve the task in the presence of background noise.

We then asked if the extrinsic input variants impacted 1) the creation and strength of multiple basins of attraction, and 2) the path of a neural trajectory towards these basins of attraction. A “strong” basin of attraction is characterized by trajectories that exhibit low-variance clusters concentrated around the basins of attraction. It is also desirable for these basins of attraction to be spatially separated, as basins that are overlapping would prevent the network from distinguishing different cue stimuli. As such, we developed a simple measure called “attractor strength coefficient” (ASC), which is calculated as 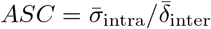 where 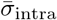 is the average intra-cluster standard deviation in PCA space for each task condition, and 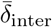 is the average distance between the median coordinates of all clusters. ASC values are bounded on the range (0,*∞*), where values closer to zero indicate the desirable properties of low intra-cluster variance, and high distances between the clusters. ASC values reported here are for trajectories limited to PC1-PC3 to match what is shown visually in Figure 7D, however we found the results hold when ASC was calculated on PC1-PC8 (8 components explains *>* 90 % variance for all network variants).

Figure 7C plots neural trajectories over time for all 4 extrinsic input variants (panels i-iv). The entire 1 second simulation is shown, with the beginning marked by a black star, and the end marked by a colored circle corresponding to each task condition. Cue inputs with NMDA_dend_ (panel i, ASC=0.066) and NMDA_soma_ (panel iii, ASC=0.056) produced a clear separation in trajectories at the start of the trial, and quickly settled into distinct regions of the state space indicating the presence of unique basins of attraction. The final positions of the trajectories produced by NMDA_soma_ (panel iii) are generally more separated than NMDA_dend_ (panel i), however trajectories for both exhibit tight clustering around their basins of attraction. In contrast, AMPA_soma_ (panel iv, ASC=0.120) produces trajectories that are clearly separable, however they visibly exhibit high variance around their basin of attraction relative to the distance between the start and end of the simulation. One possible explanation for the high variance is that persistent firing is being maintained by recurrent connections between neurons, rather than persistent bursting of individual neurons as suggested by Figure 5A(iv) and B(iv). Inspecting the learned local connectivity further supports this hypothesis, as AMPA_soma_ exhibits higher weights for all excitatory connections compared to NMDA_dend_ and NMDA_soma_ (Figure 2C(i-iii) orange histogram). A potential consequence of these weaker (high variance) basins of attraction, is sensitivity to perturbation, as sufficient noise could push the neural trajectory into a different basin causing the network to “forget” information about the cue stimulus.

The trajectories of the unsuccessfully trained AMPA_dend_(ASC=5.527) network collapse into an identical basin of attraction across all task conditions ((Figure 7C(ii))). Interestingly, the trajectories are clearly distinguishable at the start of the simulation, indicating that the failure of AMPA_dend_ to perform the working memory task is not related to its ability to produce a unique stimulus response, but instead reflects its inability to create unique and stable attractors.

Overall, analysis of PCA trajectories demonstrated that location and time constants of extrinsic synapses (and associated changes to intrinsic parameters), dramatically alter neural trajectories produced during the working memory task. A potentially important observation is that NMDA synapses at either the dendrite or soma produce strong basins of attraction (Figure 7C(i and iii)), which is supported by their relatively small ASC values (0.066 and 0.056 for NMDA_dend_ and NMDA_soma_ compared to 0.120 and 5.526 for AMPA_dend_ and AMPA_soma_). This result is not apparent from inspection of network-level spiking, as spike rasters for NMDA_dend_ (Figure 3B) are highly distinct from those of NMDA_soma_ (Figure 4B), with elevated firing activity throughout the simulation. In contrast, despite the spike rasters of NMDA_soma_ appearing more similar to AMPA_soma_ with relatively low baseline spiking activity and sharply defined clusters (Figure 4B and D), analysis of the neural trajectories potentially suggests the attractors produced by AMPA_soma_ are less stable than those produced by NMDA_soma_ (Figure 7C(iii and iv)).

We note that some of these results may depend upon the choice of using PCA for dimensionality reduction. An important flaw of analyzing neural trajectories with PCA is that the analysis does not directly account for noise or time-dependencies (dynamics), and instead treats every time point independently with the goal of capturing the covariance structure in the data [64, 66]. While these oversimplifications make PCA an imperfect fit for estimating dimensionality and state-space trajectories in neural data, PCA remains one of the most popular choices for dimensionality reduction of neuroscience, allowing this work to be more directly compared to existing publications. Further, given striking differences in PCA-estimated neural trajectories across extrinsic input variants, it is unlikely that our conclusions would change given an alternative dimensionality reduction technique.

For the goal of guiding the construction/training of biophysical network models, these results suggest attractor dynamics measured by ASC could be a good indicator of network-level activity that should be promoted during training, potentially by direct incorporation into the loss function. In contrast, PCA-estimated dimensionality did not seem to directly relate to task performance given the similarity between AMPA_dend_ and AMPA_soma_ (Figure 7A), and instead indicated a general trend of NMDA network variants exhibiting lower dimensional neural activity.

### Magnesium block supplements slow NMDA kinetics to produce stable attractors

The previous sections have demonstrated that NMDA inputs can produce stable attractors across the two different synapse locations (dendrite vs soma), whereas AMPA receptors failed to produce stable attractors through dendritic inputs. While the slow time constants of NMDA receptors are an important feature that could contribute to this difference, they also possess a magnesium ion block, which is only removed with depolarization of the postsynaptic cell [48]. This magnesium ion block is theorized to function as a coincidence detector, however its contribution to attractor dynamics is unclear.

To further identify what dynamic properties of AMPA and NMDA receptors contribute to working memory task performance, we swapped the rate constants of the NMDA and AMPA receptors. This produced two modified receptors including a fast “NMDA-like” receptor 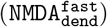, and a slow “AMPA-like” receptor 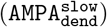. Importantly, 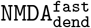 possessed fast kinetics with the characteristic magnesium block, whereas 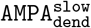 possessed slow kinetics without the magnesium block. Figure 8A (pink) demonstrates that 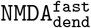 was unsuccessfully trained on the working memory task, suggesting that the magnesium block alone does not convey an advantage for performing the task. In contrast the slow kinetics of 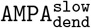 did lead to an improvement of task performance (Figure 8B, purple), however the network did not converge to the target error threshold of 1.0. Further, neural trajectory analysis of 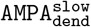 (Figure 8E) reveals no distinguishable attractor dynamics, despite the reduction in error. This result suggests that slow receptor kinetics combined with the magnesium block enable dendritic inputs to produce stable attractor dynamics, as shown in the unmodified NMDA_dend_ (Figure 8A (blue line) and Figure 7C).

**Fig 8.**
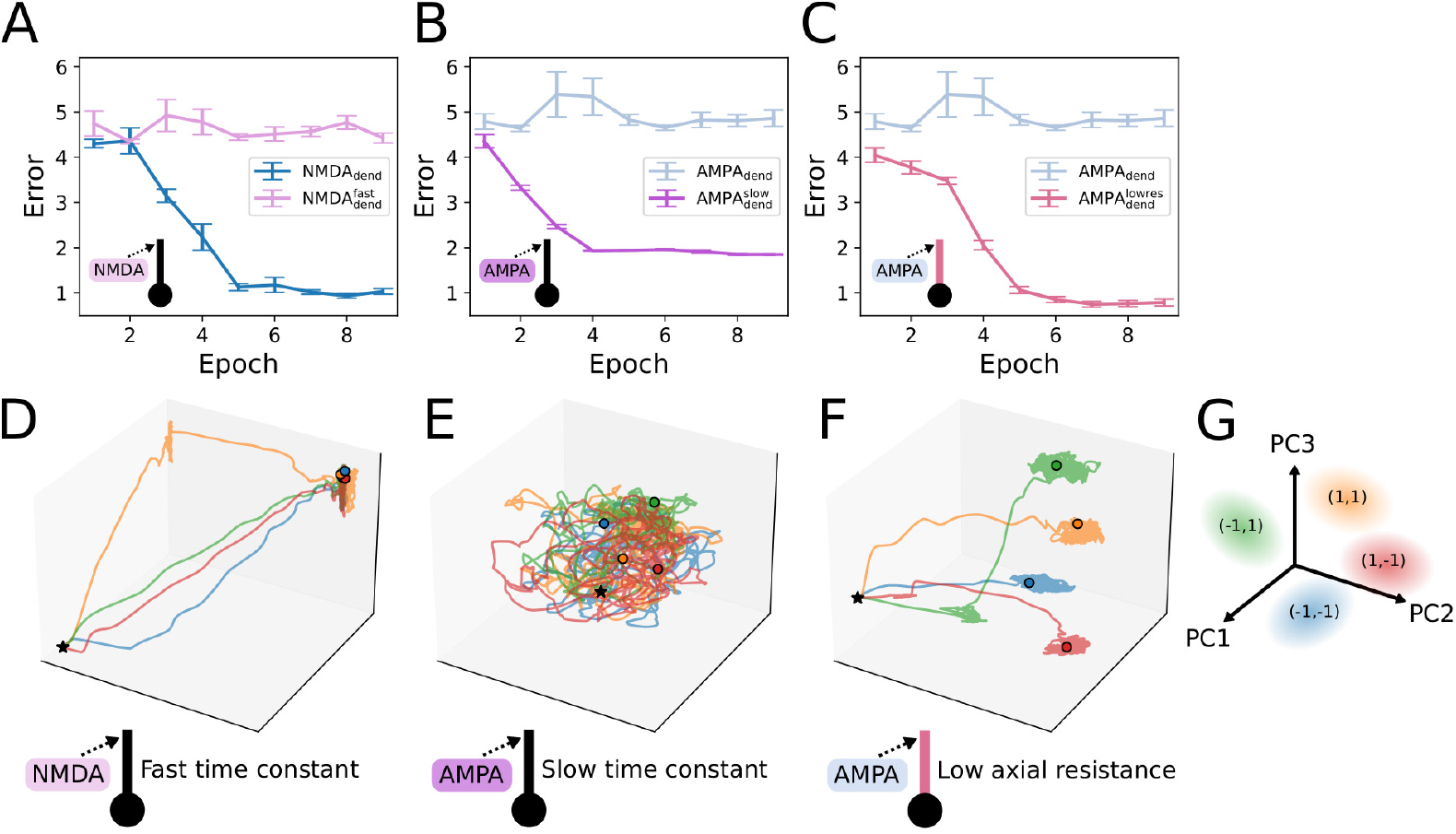
Dendritic inputs with slow rate constant or magnesium block alone do not promote attractor dynamics. **A:** NMDA receptors were modified by setting their rate constants equal to the default AMPA rate constants. This creates a fast “NMDA-like” receptor (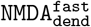, pink) that possesses a magnesium block. Compared to the successfully trained NMDA_dend_ (blue), the modified 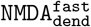 was unable to learn the working memory task. **B:** Similar modification as **A** to create a slow “AMPA-like” receptor (purple). The slow receptor kinetics did permit the network to reduce error on the working memory task, but it failed to reach the target error threshold of 1.0. The unsuccessfully trained AMPA_dend_ (light blue) is overlaid. **C** AMPA_dend_ (light blue) was retested in a network where the axial resistivity was reduced from 300Ωcm to 10Ωcm ( 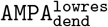, violet red). This change allowed the network to be successfully trained with dendritic AMPA receptors and reach the target error threshold. **D**: Neural trajectories associated with 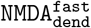 (panel A) are shown. Black stars indicate the start of the cue stimulus, colored lines indicate each task condition, and colored dots indicate the end of the 1 second simulation (same as Figure 7C). As shown, 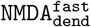 fails to produce a stable attractor, such that all trajectories collapse to the same region of the state space. **E:** Neural trajectories for 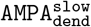 (panel B) indicate that no stable attractor is produced, and instead all trajectories overlap. **F**: Neural trajectories for 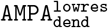 (panel C) indicate the presence of distinct and stable attractor dynamics. **G**: Legend for panels D-F labeling principal components 1-3 (axes) and task conditions (colored ovals).

### Dendritic AMPA receptors fail to produce attractor dynamics due to requirement to produce dendritic spikes

As a final investigation, we sought to identify which properties of the dendrite were responsible for the inability of AMPA_soma_to perform the working memory task. The most likely candidate is the axial resistivity, which leads to the attenuation of dendritic inputs in a purely passive dendrite. As observed in Figure 6A(i-ii), cue inputs arriving at the 3rd apical segment are propagated to the soma through dendritic action potentials. One potential issue with this constraint is that active dendrites conducive to action potentials raise the overall excitability of the neurons (e.g. increased spiking in Figure 3 compared in Figure 4). We hypothesized that AMPA_soma_ inputs are unable to overcome this increase in activity to produce stable attractor dynamics. To test if the requirement for active dendrites was indeed harming performance of dendritic AMPA inputs, we decreased the axial resistivity of the apical dendrite from 300 Ωcm to 10 Ωcm 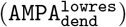. As shown in Figure 8C, this change allowed 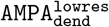 (violet red line) to rapidly reach the target error threshold, and produce distinct and persistent attractors for each task condition (Figure 8F). While further analysis will be necessary to narrow down the exact impact of active dendrites on the overall excitability of a neuron, the results clearly demonstrate that the changes in intrinsic biophysics that yield active dendrites can be harmful to task performance when task information is relayed through dendritic AMPA receptors. The unique properties of NMDA receptors (i.e. slow rate constants and magnesium block) are both essential to overcoming the associated changes in dendritic biophysics, and produce stable attractor dynamics conducive to working memory.

## Discussion

### Summary

This study introduces biophysical reservoir computing (BRC) as a framework to discover mechanistic insights that guide the construction and training of biophysically-detailed network models to solve cognitive tasks. To address the challenges of training biophysically-detailed network models, we developed a novel optimization algorithm, named “neural flow evolution”, which combines evolutionary algorithms with neural density estimators. Neural flow evolution benefits from the robust parameter search capabilities afforded by evolutionary algorithms, and the ability of neural density estimators to represent complex, high-dimensional distributions.

Training the BRC network to solve a simplified working memory task led to the following mechanistic insights with respect to extrinsic cue inputs: 1) NMDA receptors are the most effective for working memory for both somatic and dendritic synapse locations, exhibiting improved training efficiency and attractor dynamics, 2) dendritic AMPA receptors are insufficient for producing persistent spiking and associated attractor dynamics that are necessary for working memory, and 3) the combination of NMDA receptors’ unique properties, specifically their slow rate constants and magnesium block, are both required for working memory when extrinsic inputs arrive at the dendrite.

We additionally “opened the black box” [67] of the BRC, and characterized how the trained networks solved the working memory task. Since the BRC is composed of biophysically-detailed neurons, we were able to use analysis techniques common to systems neuroscience for this goal. We revealed that while NMDA receptors at the dendrite and soma are equally efficient at solving the working memory task, however, networks which delivered extrinsic cue inputs through the dendrite exhibited higher spiking activity at baseline. This difference suggests that the dendritic ion channel configurations necessary to relay cue information to the soma makes the neurons more excitable and sensitive to noisy inputs, and may explain why networks with AMPA inputs to the dendrite failed to learn the task. The spiking analysis also suggested that a synchronous burst of inhibitory neuron spiking was essential for resetting network dynamics, and subsequently activating persistent spiking in a new set of neurons which encoded the memory. We additionally revealed that single neuron burst properties were not predictive of working memory task performance, indicating that a combination of intrinsic bursting and local network connectivity underlie successfully solving the working memory task. Lastly, by examining the learned synaptic weights of the recurrent local connectivity, we revealed that the different types of extrinsic cue inputs produce highly distinct network configurations, suggesting an important role of network-level interactions in solving the working memory task.

These results are in agreement with neuroscience literature, indicating that the observed importance of NMDA receptors for working memory is not a byproduct of biological constraints, but instead reflects a real and generalizable functional benefit. Since the learned cell and network configurations in the BRC framework are not forced to match real brain activity, these results suggest that the importance of NMDA for working memory is a general computational principle that is applicable to both real and artificial neural systems.

### Related work and future directions

#### Jaxley

This study directly builds upon the software package Jaxley, which was created to directly address the challenges of training biophysical neural models by solving the complex differential equation representing biophysical models using advanced numerical computing software. These numerical computing platforms are commonly used by artificial intelligence (AI) researchers to develop deep neural networks. Specifically, Jaxley employs Jax [68], a high-performance numerical computing platform that permits auto-differentiation for backpropagation, code compilation, and parallelized computations that can be accelerated with GPU computing [27]. It is important to note that Jaxley uses standard techniques to simulate biophysical models, namely differential equation solvers like those in NEURON [69]. However, implementing these models and solvers in Jax provides the software and hardware advantages that have historically been reserved for the development and application of deep neural networks, advantages which have significantly contributed to the rapid growth of deep learning in recent years.

In [27], they train a biophysically detailed model to perform a working memory task using gradient descent with backpropagation. There are several major differences that distinguish our work from the example in this publication which we outline below:

The first major difference is that the composition of our network model was focused on neurons with active dendrites that transmitted extrinsic inputs via dendritic spikes, a feature that is suggested to convey significant computational benefits to cell and network-level function (see “Experimental studies on the computational benefits of dendrites” below). Specifically, our network model contains distinct populations of excitatory and inhibitory neurons. The excitatory neurons possess an elongated apical dendrite, a characteristic feature of cortical pyramidal neurons often implicated in top-down (feedback) information processing [70–72]. Our network model additionally uses AMPA, NMDA, and GABAa synapses for local cell-cell connections, as well AMPA and NMDA synapses for extrinsic task-relevant inputs. Mathematically, these synapses are not differentiable, preventing the use of backpropagation for gradient-based optimization of network parameters. The non-differentiability of the synapses in our model is an important limitation of our model as it prevents the use of gradient descent with backpropagation, but it permits an investigation into more biologically realistic synapses with distinct computational roles as suggested by neuroscience literature.

Instead, the network in [27] uses a differentiable synapse model for recurrent local connections that permits gradient-based optimization. The weights of these synapses were optimized such that they could produce either excitatory or inhibitory conductances, whereas our model enforces a strict separation between excitatory and inhibitory neurons. The extrinsic cue inputs in [27] were also distinct from our model, as they employed direct current injection to the soma of neurons, whereas our network model used a dedicated population of cue neurons with a gaussian receptive field that transmitted cue information with either AMPA or NMDA synapses.

The second major difference is the working memory task used. In our study we focus on a highly simplified working memory task in favor of thoroughly characterizing individual biophysical details. [27] trains their model on significantly more challenging tasks (delayed match to sample and evidence integration), to test hypotheses on different forms of neural coding.

The third major difference is our use of reservoir computing, where the parameters for individual neurons and synapses are shared across the entire network. While this potentially limits the computational capabilities of the biophysical network model, it provides a strong constraint for identifying generically useful biophysical properties. For example, our study revealed that NMDA receptors are generically useful for producing attractor dynamics, however their properties make them uniquely well-suited for transmitting information through dendrites.

It is important to emphasize that Jaxley’s ability to perform gradient-based optimization in network models with biophysically detailed neurons containing non-linear dynamics is a major advance offered by the platform, and has the potential to massively increase the complexity of tasks that biophysical models can be trained to perform. Our choice to use AMPA and NMDA synapses for the extrinsic cue input and multicompartment active pyramidal neuron dendrites was motivated by having a closer connection to neuroscience literature. An important future investigation will be to implement differentiable synapses in the BRC framework with equivalent time constants for AMPA and NMDA synapses, and a differentiable implementation of the magnesium block in the NMDA synapses. Employing differentiable synapses would allow a further investigation into the importance of NMDA for working memory as it would enable the use of gradient-based optimization, potentially expanding the complexity of working memory tasks that can be investigated. A complementary future direction will be to adapt surrogate gradient techniques from spiking neural networks [20] to perform gradient-based training in Jaxley with non-differentiable synapse models. Alternatively, FORCE training [17, 73], another method popular for task-trained RNNs and SNNs, could be adapted using Jaxley, as it does not require tracking gradients backwards through time, but instead updates model parameters mid-simulation. While non-differentiable synapses prevent backpropagation through time, FORCE training could still work with multicompartment models as it only requires gradient calculations for the current time step (i.e. recursive least squares).

### Task-trained neural models without dendrites

Prior to the introduction of Jaxley, researchers persisted in creating well-trained recurrent neural network (RNN) and spiking neural network (SNN) models of working memory using more simplified non-biophysical neuron elements. For example, researchers have developed methods to transfer learned synaptic weights of a trained RNN to a SNN [57]. Outside of working memory, researchers have successfully trained SNNs to perform a wide variety of behaviorally relevant cognitive tasks [17, 19, 74].While SNNs remain simplified relative to biophysical models, they represent a much more biologically plausible architecture compared to RNNs and modern deep learning systems.

An active area of research with RNNs is studying how a single network can be trained to perform multiple tasks [75, 76]. These studies have demonstrated that trained RNNs exhibit a form of compositionality, that is they learn dynamical motifs that are flexibly recombined to solve tasks with overlapping computational demands. One future direction of the BRC framework would be to study the emergence of compositionality in biophysical neural models trained to perform multiple tasks. A valuable point of comparison would be to train biophysical models and RNNs to perform the same array of behavioral tasks, and assess differences in how they compose dynamical motifs. Recently developed methods for comparing dynamical systems may prove instrumental for this goal [77, 78].

### Biophysical neural models with dendrites

Task-training of biophysical neural models is a relatively recent development given the high computational cost of simulation and training. Instead, biophysical neural models have been more frequently used to model specific brain regions by highly constraining model parameters and activity patterns to experimental measurements. Nevertheless, there are several biophysical neural modeling studies worth mentioning due to their implications for working memory in networks with dendrites.

With respect to biophysical models with morphologically simplified dendrites, [79] showed how at the single-neuron level, active dendrites dramatically expanded the dynamic range of neuron’s firing rate in response to synaptic inputs. In a separate study using networks of dendrite-equipped neurons, [58] demonstrated that dendritic non-nonlinearities reduced the number of interconnected neurons necessary for the flexible emergence and termination of persistent firing activity. The source of these benefits was largely linked to NMDA mediated dendritic spikes in layer 5 pyramidal neurons, and demonstrated a novel mechanism in which excitatory synaptic inputs to these neurons were able to terminate persistent activity. Building from these results, [37] showed that large networks of dendrite-equipped biophysical neurons could maintain multiple stable representations of extrinsic inputs, and that NMDA mediated persistent activity was crucial for the network to achieve multistable attractor dynamics. The key difference from our study and [37] is that they highly constrain model parameters and activity to closely match experimental measurements of real cortical circuits. Accordingly, their model agrees with the result in our study that NMDA is required under biological constraints observed in the neocortex. Our use of the BRC framework reveals a more generalized result that NMDA is generically useful for working memory, even when local connectivity and cell biophysics are freely optimized with potentially non-biological network configurations.

### Experimental studies on the computational benefits of dendrites

Unlike the neural modeling studies described previously, there are significantly fewer examples of experimental studies investigating the role of dendrites in working memory. However, numerous experimental studies have proposed a diverse set of alternative computational benefits that are unique to dendrites. These proposals include computational benefits such as compartmentalization [80], coincidence detection [81, 82], modulation of sensory perception [83], and learning [84].

Despite the fact that all of these theories concern neurons in a highly interconnected network, the computational benefits of these theories primarily concern functions at the level of single neurons (see [85]). Experimental studies examining the relationship between dendrites and their impact on network level dynamics are significantly fewer in number due to the technical challenges of performing such experiments [26]. In future work, the BRC framework may prove to be a highly useful tool for isolating the computational benefits of dendrites beyond the simple working memory task presented here, and propose new experiments that take into account the network-level functioning of dendrite-equipped neurons.

### Limitations

There are several limitations to the present study that provide ample opportunity for follow up research.

The first limitation is the simple morphology used for the apical dendrite. As stated in the introduction, a primary motivation of this study was understanding which modeling choices made sense when expanding task-trained RNNs and SNNs to include dendrites. Our results revealed the dendritic NMDA receptors are uniquely suited for transmitting extrinsic cue information, and ultimately recruit distinct clusters of persistently firing neurons. However, we did not observe a unique benefit of transmitting cue information through dendrites, and instead demonstrated the dendritic AMPA receptors were insufficient to promote attractor dynamics. It is important to emphasize that these results do not suggest dendrites are not computationally useful for other tasks. Many of the theories of dendritic function emphasize highly elaborated dendritic arbors which serve to process incoming information separately from the soma. A valuable expansion of the BRC framework would be the inclusion of morphologically complex dendrites to directly test more sophisticated theories of dendritic function.

The second limitation is the simplicity of the cognitive task used. We focused on a basic working memory which required maintaining persistent representations of 4 distinct stimuli. In general, the cognitive tasks used in experimental settings are highly simplified with respect to the behavioral demands of the real world [86–88]. While our simplified working memory task is not representative of the cognitive demands of naturalistic behavior, we argue that persistent representations are a fundamental building block required for a vast array of more complex functions. Further, the simplified working memory task was surprisingly sufficient to demonstrate the fundamental role of NMDA receptors in the context of persistent representations. It is highly possible that the network implemented will be insufficient for more complex cognitive tasks, which may require additional biophysical properties, optimization strategies, and a significantly better understanding of how these computations are performed in real neural circuits.

The third limitation is the exclusive use of AMPA and GABAa receptors for recurrent local connectivity. This choice was made early in the study, as inclusion of recurrent NMDA synapses was found to produce runaway excitation, a problem that may reflect the relatively small number of neurons in the biophysical reservoir. However, experimental studies have demonstrated that NMDA and AMPA receptors are both involved in the onset and maintenance of persistent firing underlying cortical working memory [35, 36]). Follow up studies will be necessary to elucidate the role of recurrent local NMDA connections, and if their inclusion can mitigate the issues we report here with signaling cue information through dendritic AMPA synapses.

The fourth limitation is the lack of heterogeneity in cell-type parameters of the network. It is well documented that cell heterogeneity plays a crucial role in the computational properties of biologically-plausible neural networks [89–92]. However, in the context of working memory, [90] demonstrates with SNNs that cell heterogeneity (e.g. varied spike thresholds) actually impairs the ability of networks to encode and maintain information about external stimuli. Cell heterogeneity was shown to increase the dimensionality of neural activity, making these networks more suitable as function generators that continuously transform inputs to a new target signal. A future direction will be to explore if this tradeoff between function generation and persistent representations holds in network models composed of biophysically detailed neurons, and if heterogeneity in dendritic parameters confer unique computational properties. One possibility is that the presence of connection heterogeneity (i.e. between cell connection probabilities) in biophysically detailed neurons with dendritic synapses provides similar benefits as cell heterogeneity in SNNs.

As described above, the decision to use evolutionary algorithms combined with neural density estimation (see “Neural flow evolution” in Methods) was largely due to challenges we faced with existing optimization techniques. Previous work, including our own, has shown that neural density estimation is particularly effective for parameter estimation in biophysical models [56, 93]. Traditionally this takes the form of likelihood-free Bayesian inference, where the estimator learns the posterior distribution *p*(*θ | x*) over model parameters *θ* conditioned on simulated outputs *x*. While these methods have been effective for fitting biophysical models to data, we found that conditional density estimation was numerically unstable in the setting of training the BRC to solve the working memory, with the majority of parameter samples falling outside the support of the prior distribution. Instead, we used neural density estimation in a simpler form to approximate an unconditional distribution *p*(*θ*). We note that previous studies have proposed integrating probability distributions with evolutionary algorithms as a method to generate offspring from the parent population [94]. To our knowledge, this is the first study to propose using neural density estimation for this purpose. Follow up work will be necessary to benchmark this approach against standard evolutionary algorithms, as well as traditional density estimation techniques like kernel density estimation.

## Code availability

All code used for simulation and analysis in this manuscript is located at https://github.com/ntolley/dendractor

## Supplemental Data

## Acknowledgments

This work was supported by the Brown Biomedical Innovation to Impact Award, the National Institute of Health (NIH; https://www.nih.gov; grant number U24NS129945), and the National Science Foundation (NSF; https://www.nsf.gov; grant number 2424101). The funders had no role in study design, data collection and analysis, decision to publish, or preparation of the manuscript.

We would like to thank Dr. Katharina Duecker, Dr. Jason Ritt, and Dr. Christopher Moore for feedback on early versions of this manuscript.

## References

1. Yu Y, Si X, Hu C, Zhang J. A Review of Recurrent Neural Networks: LSTM Cells and Network Architectures;31(7):1235–1270. doi:10.1162/necoa01199.

2. Soo WWM, Goudar V, Wang XJ. Training biologically plausible recurrent neural networks on cognitive tasks with long-term dependencies;. Available from: https://www.biorxiv.org/content/10.1101/2023.10.10.561588v1.

3. Pals M, Macke JH, Barak O. Trained recurrent neural networks develop phase-locked limit cycles in a working memory task;20(2):e1011852. doi:10.1371/journal.pcbi.1011852.

4. Vyas S, Golub MD, Sussillo D, Shenoy KV. Computation Through Neural Population Dynamics;43:249–275. doi:10.1146/annurev-neuro-092619-094115.

5. Pascanu R, Jaeger H. A neurodynamical model for working memory;24(2):199–207. doi:10.1016/j.neunet.2010.10.003.

6. Yang GR, Molano-Mazón M. Towards the next generation of recurrent network models for cognitive neuroscience;70:182–192. doi:10.1016/j.conb.2021.10.015.

7. Khan A, Sohail A, Zahoora U, Qureshi AS. A survey of the recent architectures of deep convolutional neural networks;53(8):5455–5516. doi:10.1007/s10462-020-09825-6.

8. Taylor L, Nitschke G. Improving Deep Learning with Generic Data Augmentation. In: 2018 IEEE Symposium Series on Computational Intelligence (SSCI);. p. 1542–1547. Available from: https://ieeexplore.ieee.org/abstract/document/8628742.

9. Shorten C, Khoshgoftaar TM. A survey on Image Data Augmentation for Deep Learning;6(1):60. doi:10.1186/s40537-019-0197-0.

10. Narkhede MV, Bartakke PP, Sutaone MS. A review on weight initialization strategies for neural networks;55(1):291–322. doi:10.1007/s10462-021-10033-z.

11. Hanin B, Rolnick D. How to Start Training: The Effect of Initialization and Architecture. In: Advances in Neural Information Processing Systems. vol. 31. Curran Associates, Inc.;.Available from: https://proceedings.neurips.cc/paper/2018/hash/d81f9c1be2e08964bf9f24b15f0e4900-Abstract.html.

12. Salimans T, Kingma DP. Weight Normalization: A Simple Reparameterization to Accelerate Training of Deep Neural Networks. In: Advances in Neural Information Processing Systems. vol. 29. Curran Associates, Inc.;.Available from: https://proceedings.neurips.cc/paper/2016/hash/ed265bc903a5a097f61d3ec064d96d2e-Abstract.html.

13. Janocha K, Czarnecki WM. On Loss Functions for Deep Neural Networks in Classification;. Available from: http://arxiv.org/abs/1702.05659.

14. Wang Q, Ma Y, Zhao K, Tian Y. A Comprehensive Survey of Loss Functions in Machine Learning;9(2):187–212. doi:10.1007/s40745-020-00253-5.

15. Eshraghian JK, Ward M, Neftci EO, Wang X, Lenz G, Dwivedi G, et al. Training spiking neural networks using lessons from deep learning;111(9):1016–1054.

16. Kim R, Li Y, Sejnowski TJ. Simple framework for constructing functional spiking recurrent neural networks;116(45):22811–22820. doi:10.1073/pnas.1905926116.

17. Nicola W, Clopath C. Supervised learning in spiking neural networks with FORCE training;8(1):2208. doi:10.1038/s41467-017-01827-3.

18. Arthur BJ, Kim CM, Chen S, Preibisch S, Darshan R. A scalable implementation of the recursive least-squares algorithm for training spiking neural networks;17. doi:10.3389/fninf.2023.1099510.

19. Wang G, Sun Y, Cheng S, Song S. Evolving Connectivity for Recurrent Spiking Neural Networks;36:2991–3007.

20. Neftci EO, Mostafa H, Zenke F. Surrogate Gradient Learning in Spiking Neural Networks: Bringing the Power of Gradient-Based Optimization to Spiking Neural Networks;36(6):51–63. doi:10.1109/MSP.2019.2931595.

21. D’Esposito M. From cognitive to neural models of working memory;362(1481):761–772. doi:10.1098/rstb.2007.2086.

22. Murray JD, Jaramillo J, Wang XJ. Working Memory and Decision-Making in a Frontoparietal Circuit Model;37(50):12167–12186. doi:10.1523/JNEUROSCI.0343-17.2017.

23. Gilhooly KJ. Working memory and planning. In: The Cognitive Psychology of Planning. Psychology Press;.

24. Rhodes BJ, Bullock D, Verwey WB, Averbeck BB, Page MPA. Learning and production of movement sequences: Behavioral, neurophysiological, and modeling perspectives;23(5):699–746. doi:10.1016/j.humov.2004.10.008.

25. Larkum ME. Are Dendrites Conceptually Useful?;489:4–14. doi:10.1016/j.neuroscience.2022.03.008.

26. Poirazi P, Papoutsi A. Illuminating dendritic function with computational models;21(6):303–321. doi:10.1038/s41583-020-0301-7.

27. Deistler M, Kadhim KL, Pals M, Beck J, Huang Z, Gloeckler M, et al. Differentiable simulation enables large-scale training of detailed biophysical models of neural dynamics;. Available from: https://www.biorxiv.org/content/10.1101/2024.08.21.608979v1.

28. Tanaka G, Yamane T, Héroux JB, Nakane R, Kanazawa N, Takeda S, et al. Recent advances in physical reservoir computing: A review;115:100–123.

29. Damicelli F, Hilgetag CC, Goulas A. Brain connectivity meets reservoir computing;18(11):e1010639. doi:10.1371/journal.pcbi.1010639.

30. Zhang H, Vargas DV. A Survey on Reservoir Computing and its Interdisciplinary Applications Beyond Traditional Machine Learning;11:81033–81070. doi:10.1109/ACCESS.2023.3299296.

31. Hasson U, Nastase SA, Goldstein A. Direct Fit to Nature: An Evolutionary Perspective on Biological and Artificial Neural Networks;105(3):416–434. doi:10.1016/j.neuron.2019.12.002.

32. Erhan D, Courville A, Bengio Y, Vincent P. Why does unsupervised pre-training help deep learning? JMLR Workshop and Conference Proceedings;. p. 201–208.

33. Larochelle H, Bengio Y, Louradour J, Lamblin P. Exploring strategies for training deep neural networks.;10(1).

34. Wang XJ. 50 years of mnemonic persistent activity: quo vadis?;44(11):888–902. doi:10.1016/j.tins.2021.09.001.

35. Self MW, Kooijmans RN, Supèr H, Lamme VA, Roelfsema PR. Different glutamate receptors convey feedforward and recurrent processing in macaque V1;109(27):11031–11036. doi:10.1073/pnas.1119527109.

36. Vugt Bv, Kerkoerle Tv, Vartak D, Roelfsema PR. The Contribution of AMPA and NMDA Receptors to Persistent Firing in the Dorsolateral Prefrontal Cortex in Working Memory;40(12):2458–2470. doi:10.1523/JNEUROSCI.2121-19.2020.

37. Stefanos SS, Papoutsi A, Poirazi P. Structured connectivity exploits NMDA non-linearities to enable flexible encoding of multiple memoranda in a PFC circuit model;. Available from: https://www.biorxiv.org/content/10.1101/733519v2.

38. Wang M, Yang Y, Wang CJ, Gamo NJ, Jin LE, Mazer JA, et al. NMDA Receptors Subserve Persistent Neuronal Firing during Working Memory in Dorsolateral Prefrontal Cortex;77(4):736–749. doi:10.1016/j.neuron.2012.12.032.

39. Abbott LF, Dayan P. Theoretical Neuroscience: Computational and Mathematical Modeling of Neural Systems. Computational Neuroscience Series. MIT Press;.

40. Bittner KC, Andrasfalvy BK, Magee JC. Ion Channel Gradients in the Apical Tuft Region of CA1 Pyramidal Neurons;7(10):e46652. doi:10.1371/journal.pone.0046652.

41. Rhodes P. The Properties and Implications of NMDA Spikes in Neocortical Pyramidal Cells;26(25):6704–6715. doi:10.1523/JNEUROSCI.3791-05.2006.

42. Palmer LM, Shai AS, Reeve JE, Anderson HL, Paulsen O, Larkum ME. NMDA spikes enhance action potential generation during sensory input;17(3):383–390. doi:10.1038/nn.3646.

43. Schulz JM, Knoflach F, Hernandez MC, Bischofberger J. Dendrite-targeting interneurons control synaptic NMDA-receptor activation via nonlinear 5-GABAA receptors;9(1):3576. doi:10.1038/s41467-018-06004-8.

44. Olàh VJ, Wu J, Kaczmarek LK, Rowan MJ. Input-specific gating of NMDA amplification via HCN channels in mouse L2/3 pyramidal neurons;13. doi:10.7554/eLife.96002.2.

45. Major G, Larkum ME, Schiller J. Active Properties of Neocortical Pyramidal Neuron Dendrites;36:1–24. doi:10.1146/annurev-neuro-062111-150343.

46. Pagkalos M, Chavlis S, Poirazi P. Introducing the Dendrify framework for incorporating dendrites to spiking neural networks;14(1):131. doi:10.1038/s41467-022-35747-8.

47. Pospischil M, Toledo-Rodriguez M, Monier C, Piwkowska Z, Bal T, Frégnac Y, et al. Minimal Hodgkin–Huxley type models for different classes of cortical and thalamic neurons;99:427–441.

48. Destexhe A, Mainen ZF, Sejnowski TJ. Kinetic models of synaptic transmission;2:1–25.

49. Watts DJ, Strogatz SH. Collective dynamics of ‘small-world’ networks;393(6684):440–442. doi:10.1038/30918.

50. Sporns O, Zwi JD. The small world of the cerebral cortex;2(2):145–162. doi:10.1385/NI:2:2:145.

51. Shanahan M. Dynamical complexity in small-world networks of spiking neurons;78(4):041924. doi:10.1103/PhysRevE.78.041924.

52. Pazzini R, Kinouchi O, Costa AA. Neuronal avalanches in Watts-Strogatz networks of stochastic spiking neurons;104(1):014137. doi:10.1103/PhysRevE.104.014137.

53. Yu X, Gen M. Introduction to evolutionary algorithms. Springer Science & Business Media;.

54. Papamakarios G. Neural Density Estimation and Likelihood-free Inference;. Available from: http://arxiv.org/abs/1910.13233.

55. Pastore M, Calcagnì A. Measuring Distribution Similarities Between Samples: A Distribution-Free Overlapping Index;10. doi:10.3389/fpsyg.2019.01089.

56. Tolley N, Rodrigues PLC, Gramfort A, Jones SR. Methods and considerations for estimating parameters in biophysically detailed neural models with simulation based inference;20(2):e1011108. doi:10.1371/journal.pcbi.1011108.

57. Kim R, Sejnowski TJ. Strong inhibitory signaling underlies stable temporal dynamics and working memory in spiking neural networks;24(1):129–139. doi:10.1038/s41593-020-00753-w.

58. Papoutsi A, Sidiropoulou K, Poirazi P. Dendritic Nonlinearities Reduce Network Size Requirements and Mediate ON and OFF States of Persistent Activity in a PFC Microcircuit Model;10(7):e1003764. doi:10.1371/journal.pcbi.1003764.

59. Gallego JA, Perich MG, Miller LE, Solla SA. Neural Manifolds for the Control of Movement;94(5):978–984. doi:10.1016/j.neuron.2017.05.025.

60. Langdon C, Engel TA. Latent circuit inference from heterogeneous neural responses during cognitive tasks;28(3):665–675. doi:10.1038/s41593-025-01869-7.

61. Compte A. Computational and in vitro studies of persistent activity: Edging towards cellular and synaptic mechanisms of working memory;139(1):135–151. doi:10.1016/j.neuroscience.2005.06.011.

62. Jazayeri M, Ostojic S. Interpreting neural computations by examining intrinsic and embedding dimensionality of neural activity;70:113–120. doi:10.1016/j.conb.2021.08.002.

63. Altan E, Solla SA, Miller LE, Perreault EJ. Estimating the dimensionality of the manifold underlying multi-electrode neural recordings;17(11):e1008591. doi:10.1371/journal.pcbi.1008591.

64. Cunningham JP, Yu BM. Dimensionality reduction for large-scale neural recordings;17(11):1500–1509. doi:10.1038/nn.3776.

65. Badre D, Bhandari A, Keglovits H, Kikumoto A. The dimensionality of neural representations for control;38:20–28. doi:10.1016/j.cobeha.2020.07.002.

66. Pellegrino A, Stein H, Cayco-Gajic NA. Dimensionality reduction beyond neural subspaces with slice tensor component analysis;27(6):1199–1210. doi:10.1038/s41593-024-01626-2.

67. Sussillo D, Barak O. Opening the Black Box: Low-Dimensional Dynamics in High-Dimensional Recurrent Neural Networks;25(3):626–649. doi:10.1162/NECOa00409.

68. Bradbury J, Frostig R, Hawkins P, Johnson MJ, Leary C, Maclaurin D, et al. JAX: Autograd and XLA; p. ascl:2111.002.

69. Hines ML, Carnevale NT. The NEURON Simulation Environment;9(6):1179–1209. doi:10.1162/neco.1997.9.6.1179.

70. Harris KD, Shepherd GMG. The neocortical circuit: themes and variations;18(2):170–181. doi:10.1038/nn.3917.

71. Schuman B, Dellal S, Prönneke A, Machold R, Rudy B. Neocortical Layer 1: An Elegant Solution to Top-Down and Bottom-Up Integration;44:221–252. doi:10.1146/annurev-neuro-100520-012117.

72. Fişek M, Herrmann D, Egea-Weiss A, Cloves M, Bauer L, Lee TY, et al. Cortico-cortical feedback engages active dendrites in visual cortex;617(7962):769–776. doi:10.1038/s41586-023-06007-6.

73. Sussillo D, Abbott LF. Generating Coherent Patterns of Activity from Chaotic Neural Networks;63(4):544–557. doi:10.1016/j.neuron.2009.07.018.

74. Maes A, Barahona M, Clopath C. Learning spatiotemporal signals using a recurrent spiking network that discretizes time;16(1):e1007606. doi:10.1371/journal.pcbi.1007606.

75. Driscoll LN, Shenoy K, Sussillo D. Flexible multitask computation in recurrent networks utilizes shared dynamical motifs;27(7):1349–1363. doi:10.1038/s41593-024-01668-6.

76. Yang GR, Joglekar MR, Song HF, Newsome WT, Wang XJ. Task representations in neural networks trained to perform many cognitive tasks;22(2):297–306. doi:10.1038/s41593-018-0310-2.

77. Ostrow M, Eisen A, Kozachkov L, Fiete I. Beyond Geometry: Comparing the Temporal Structure of Computation in Neural Circuits with Dynamical Similarity Analysis;. Available from: http://arxiv.org/abs/2306.10168.

78. Guilhot Q, Wójcik M, Achterberg J, Costa RP. Dynamical similarity analysis can identify compositional dynamics developing in RNNs;. Available from: http://arxiv.org/abs/2410.24070.

79. Gollo LL, Kinouchi O, Copelli M. Active Dendrites Enhance Neuronal Dynamic Range;5(6):e1000402. doi:10.1371/journal.pcbi.1000402.

80. Polsky A, Mel BW, Schiller J. Computational subunits in thin dendrites of pyramidal cells;7(6):621–627. doi:10.1038/nn1253.

81. Larkum ME, Zhu JJ, Sakmann B. A new cellular mechanism for coupling inputs arriving at different cortical layers;398(6725):338–341. doi:10.1038/18686.

82. Xu Nl, Harnett MT, Williams SR, Huber D, O’Connor DH, Svoboda K, et al. Nonlinear dendritic integration of sensory and motor input during an active sensing task;492(7428):247–251. doi:10.1038/nature11601.

83. Takahashi N, Oertner TG, Hegemann P, Larkum ME. Active cortical dendrites modulate perception;354(6319):1587–1590. doi:10.1126/science.aah6066.

84. McBride TJ, Rodriguez-Contreras A, Trinh A, Bailey R, DeBello WM. Learning Drives Differential Clustering of Axodendritic Contacts in the Barn Owl Auditory System;28(27):6960–6973. doi:10.1523/JNEUROSCI.1352-08.2008.

85. Jones IS, Kording KP. Might a Single Neuron Solve Interesting Machine Learning Problems Through Successive Computations on Its Dendritic Tree?;33(6):1554–1571. doi:10.1162/necoa01390.

86. Huk A, Bonnen K, He BJ. Beyond Trial-Based Paradigms: Continuous Behavior, Ongoing Neural Activity, and Natural Stimuli;38(35):7551–7558. doi:10.1523/JNEUROSCI.1920-17.2018.

87. Maselli A, Gordon J, Eluchans M, Lancia GL, Thiery T, Moretti R, et al. Beyond simple laboratory studies: Developing sophisticated models to study rich behavior;46:220–244. doi:10.1016/j.plrev.2023.07.006.

88. Krakauer JW, Ghazanfar AA, Gomez-Marin A, MacIver MA, Poeppel D. Neuroscience Needs Behavior: Correcting a Reductionist Bias;93(3):480–490. doi:10.1016/j.neuron.2016.12.041.

89. Perez-Nieves N, Leung VCH, Dragotti PL, Goodman DFM. Neural heterogeneity promotes robust learning;12(1):5791. doi:10.1038/s41467-021-26022-3.

90. Gast R, Solla SA, Kennedy A. Neural heterogeneity controls computations in spiking neural networks;121(3):e2311885121. doi:10.1073/pnas.2311885121.

91. Lengler J, Jug F, Steger A. Reliable Neuronal Systems: The Importance of Heterogeneity;8(12):e80694. doi:10.1371/journal.pone.0080694.

92. Mejias JF, Longtin A. Differential effects of excitatory and inhibitory heterogeneity on the gain and asynchronous state of sparse cortical networks;8. doi:10.3389/fncom.2014.00107.

93. Gonçalves PJ, Lueckmann JM, Deistler M, Nonnenmacher M, Ö cal K, Bassetto G, et al. Training deep neural density estimators to identify mechanistic models of neural dynamics;9:e56261. doi:10.7554/eLife.56261.

94. Kern S, Müller SD, Hansen N, Büche D, Ocenasek J, Koumoutsakos P. Learning probability distributions in continuous evolutionary algorithms – a comparative review;3(1):77–112. doi:10.1023/B:NACO.0000023416.59689.4e.

